# Mechanism of glycogen synthase inactivation and interaction with glycogenin

**DOI:** 10.1101/2021.11.14.468517

**Authors:** Laura Marr, Dipsikha Biswas, Leonard A. Daly, Christopher Browning, Sarah Vial, Daniel P. Maskell, Catherine Hudson, John Pollard, Jay Bertrand, Neil A. Ranson, Heena Khatter, Claire E. Eyers, Kei Sakamoto, Elton Zeqiraj

## Abstract

Glycogen is the major glucose reserve in eukaryotes, and defects in glycogen metabolism and structure lead to disease. Glycogenesis involves interaction of glycogenin (GN) with glycogen synthase (GS), where GS is activated by glucose-6-phosphate (G6P) and inactivated by phosphorylation. We describe the 2.6 Å resolution cryo-EM structure of phosphorylated human GS revealing an autoinhibited GS tetramer flanked by two GN dimers. Phosphorylated N- and C-termini from two GS protomers converge near the G6P-binding pocket and buttress against GS regulatory helices. This keeps GS in an inactive conformation mediated by phospho-Ser641 interactions with a composite “arginine cradle”. Structure-guided mutagenesis perturbing interactions with phosphorylated tails led to increased basal/unstimulated GS activity. We propose that multivalent phosphorylation supports GS autoinhibition through interactions from a dynamic “spike” region, allowing a tuneable rheostat for regulating GS activity. This work therefore provides new insights into glycogen synthesis regulation and facilitates studies of glycogen-related diseases.

## Introduction

Glycogen is a branched polymer of glucose that functions as the primary energy store in eukaryotes. In its mature form, the glycogen particle can comprise up to ~50,000 glucose units that are rapidly utilized when glucose levels are low. Glycogen is stored predominantly in the muscle and liver cells, and to a lesser extent in other organs and tissues including kidney, brain, fat and heart^1^.

Glycogen is synthesized through the cooperative action of three enzymes: glycogenin (GN), glycogen synthase (GS) and glycogen branching enzyme (GBE)^2^. GN initiates the process via auto-glucosylation of a conserved tyrosine residue, producing a primer glucose chain of 8-12 resides connected by α-1,4-linkages^3^ (**Fig. 1a**). This glycogen initiating particle is further extended by GS after its recruitment by the GN C-terminus allowing the addition of glucose residues using α-1,4-linkages^4,5^. GBE introduces α-1,6-linkages every 6-8 residues to the growing glycogen molecule, thus creating the final globular structure containing GN at the centre^2,6^ (**Fig. 1b**). Glycogen exists as a population of molecules with varying sizes (10-290 nm) in different tissues and species, although the importance of this variability is not well understood^1,7^.

**Fig. 1.**
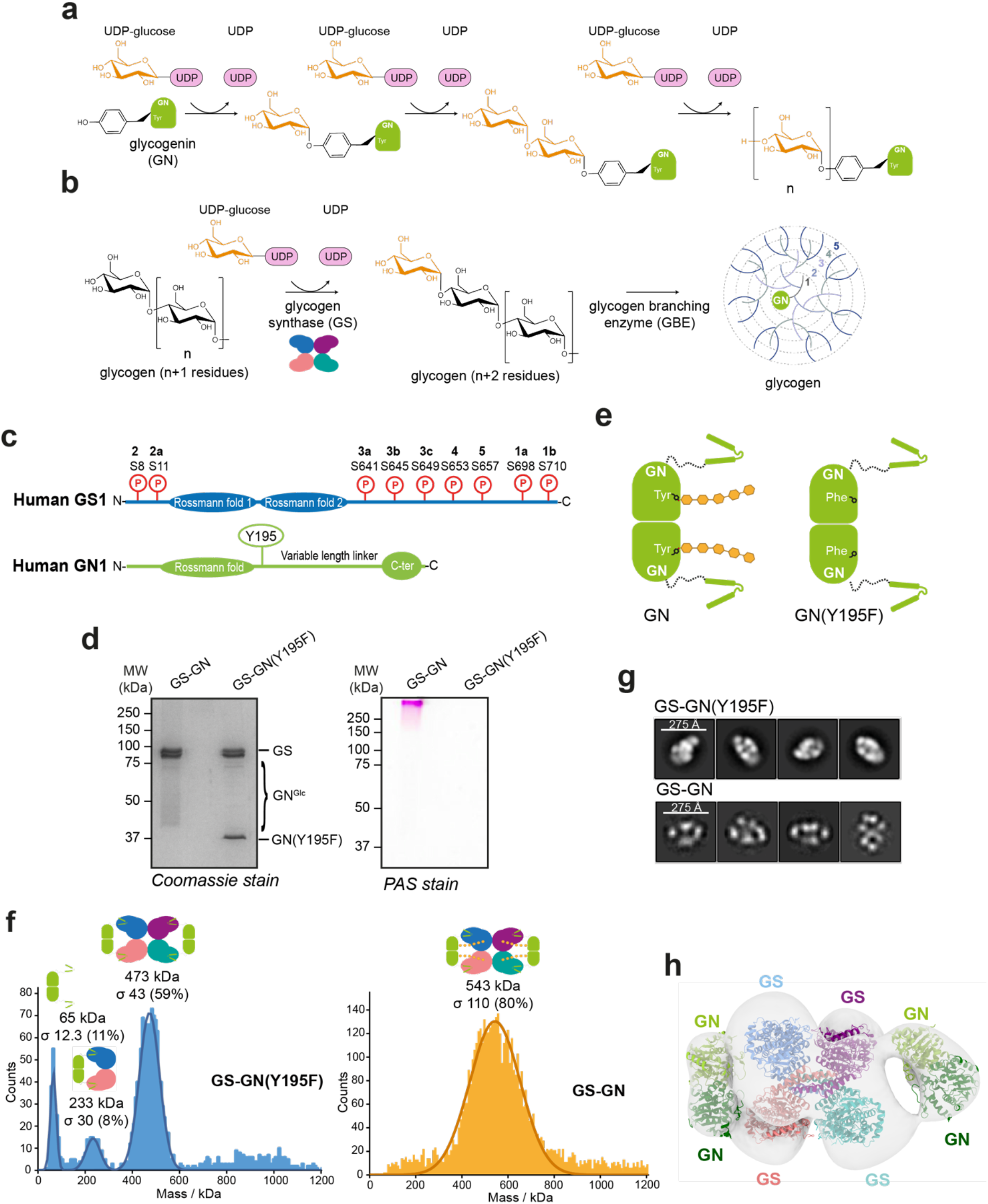
Structural analysis of the full-length GS-GN complex. **a** Enzymatic reaction catalyzed by GN. **b** Enzymatic reaction catalyzed by GS and subsequent branching of glycogen by GBE. **c** Domain architecture of human GS (top) and GN (bottom). Known *in vivo* phosphorylation sites of GS are shown in red and are labelled with residue number and classical nomenclature (in bold). GN tyrosine 195 that becomes auto-glucosylated and mutated to a phenylalanine (Y195F) in this study is indicated. Not to scale. **d** SDS-PAGE analysis of GS-GN WT and Y195F complexes (left) and periodic acid-Schiff (PAS) staining of both complexes (right). **e** Cartoon representation of GN WT and Y195F. **f** Mass photometry of GS-GN(Y195F) (left) and WT complex (right). Expected stoichiometry for each peak is indicated. The percentage of particles contributing to each peak is shown in brackets. **g** Selected 2D class averages after negative-stain electron microscopy (nsEM) analysis of indicated GS-GN complexes. **h** nsEM final map (C1 symmetry at ~22 Å) is shown in transparent surface, with fitted human GN crystal structure (PDB ID 3T7O) and human GS cryo-EM structure (reported here).

Glycogen synthesis and breakdown are tightly regulated processes, and thus dysregulation of the enzymes involved in glycogen metabolism contributes to glycogen storage diseases (GSDs), diabetes, neuroinflammation, neurodegeneration and muscle damage^1,8^. Excessive and/or abnormal glycogen is a common characteristic in most GSDs. Pompe disease (GSDII) is caused by deficiency of acid-a-glucosidase, resulting in accumulation of lysosomal glycogen and consequent lysosomal destruction and dysfunction^9^. Lafora disease is a fatal neurodegenerative condition, characterized by Lafora bodies that contain hyperphosphorylated and poorly branched, insoluble glycogen deposits^10^. In addition, loss of GS-GN interaction results in muscle weakness and cardiomyopathy^11^.

Studies using mouse models have found inhibition of glycogen synthesis, particularly by reducing GS activity, to be beneficial for multiple GSDs^12–16^. To date there is no structure of the GS-GN complex and no structure of human GS. Since inhibition of GS activity is potentially beneficial for GSD patients, obtaining a human GS-GN structure and understanding how GS is regulated is instrumental in developing new therapeutics.

GN is found in two isoforms, GN1 and GN2, encoded by the *GYG1* and *GYG2* genes respectively. While *GYG1* is widely expressed, *GYG2* is restricted to the liver, pancreas and heart^17,18^. GN belongs to the GT8 family of glycosyltransferases, containing a glycosyl transferase A (GT-A) fold with a single Rossmann fold domain at the N-terminus, which is essential for binding of the glucose donor uridine diphosphate glucose (UDP-G)^19–21^. The C-terminus comprises a highly conserved region of ~34 residues (GN^34^) which is the minimal targeting region for binding GS^5,22^. Other interaction interfaces have been suggested^23^, but, further investigation into the full-length complex is required to precisely define any additional interaction interfaces. The area between the N-terminal catalytic domain and C-terminal GS binding motif is a linker region that is variable in sequence and in length (**Fig. 1c and Supplementary Fig. 1**).

GS is also found as two isoforms, GS1 and GS2, encoded by the *GYS1* and *GYS2* genes respectively. These are differentially expressed, with *GYS1* being expressed predominantly in skeletal muscle and most other cell types where glycogen is present, while *GYS2* is expressed exclusively in the liver^24–26^. Eukaryotic GS belongs to the GT3 family of glycosyltransferases with a GT-B architecture comprising an N-terminal and a C-terminal Rossmann fold domain, with an interdomain cleft that contains the active site^19,27^. GS is the rate limiting enzyme in glycogen biosynthesis and as such its activity is tightly regulated^28^. GS is inactivated by covalent phosphorylation at numerous N- and C-terminal sites (**Fig. 1c**), and is allosterically activated by glucose-6-phosphate (G6P) binding and/or dephosphorylation^2,29,30^. Human GS phosphorylation sites lie at the N-terminus (sites 2 and 2a) and C-terminus (sites 3a, 3b, 3c, 4, 5, 1a, 1b), and phosphorylation occurs in a hierarchical fashion, whereby the phosphorylation of a specific site is the recognition motif for subsequent phosphorylation^31–33^ (**Fig. 1c and Supplementary Fig. 2**). How the metazoan GS is inhibited is not clear and while allosteric activation by G6P binding been described for the yeast GS paralogues^34^ no structural information of the phosphorylated version of the enzyme exists.

The complex interplay between allosteric activation and inhibitory phosphorylation is not yet fully understood, at least in part because of the lack of structural data for the full GS-GN complex. Although a binary GS-GN complex was co-purified over 30 years ago^3^, we have yet to confirm the stoichiometry of this complex and identify precisely how the two proteins cooperate to make glycogen.

Here, we report the structural and functional analysis of the full-length human GS-GN complex and the cryo-EM structure of phosphorylated human GS. The structure reveals that phosphoregulatory elements form a flexible inter-subunit “spike” region emanating from two GS protomers, which help to keep GS in an inactive conformation via interactions of phosphorylated Ser641 (site 3a) with arginine residues from GS regulatory helices, which we have termed the arginine cradle. Moreover, low resolution maps of GN bound to GS reveal two flexible GN dimers coordinating a GS tetramer, providing new insights into the stoichiometry and the conformational plasticity of this enzyme complex. Collectively, these results shed light on the regulation of glycogen biosynthesis and the inner workings of how GS and GN cooperate to synthesize glycogen.

## Results

### GS-GN forms an equimolar 4:4 complex

To characterize the synthesis of glycogen by the GS-GN complex, we expressed and purified human full length GS1 and GN1 in insect cells. Consistent with previous reports, co-expression of GS with GN resulted in improved production yields over the expression of GS alone^35,36^. Purification of the wild-type (WT) complex resulted in a highly glucosylated sample, as evidenced by a smear by SDS-PAGE corresponding to glucosylated GN detected by Coomassie stain, periodic acid-Schiff (PAS) staining and immunoblotting (**Fig. 1d and Supplementary Fig. 3b and 3c**). In-gel protease digestion of different molecular weight regions (encompassing mass ranges from 43-55 kDa, 55-72 kDa, 95-130 kDa and greater than 130 kDa) combined with tandem mass spectrometry confirmed the presence of GN1 in all these higher MW species (**Supplementary Data 1**). In addition, treatment of GS-GN preparations with α-amylase (endo-α-1,4-d-glucan hydrolase) resulted in the disappearance of the smeared bands revealing a single, sharp band migrating at the expected molecular weight for GN1 (~37.5 kDa) and also absence of glucosylated species after PAS staining. Thus, confirming that the smearing effect is due to glucosylation of GN (**Supplementary Fig. 3d**). Mutation of the GN auto-glucosylating tyrosine 195^37,38^ to a phenylalanine (Y195F), resulted in a non-glucosylated GN species, as shown by a single band for GN migrating at the expected size (~37.5 kDa) detected by Coomassie stain and immunoblotting and absence of glucosylated species after PAS staining (**Fig. 1e, Supplementary Fig. 3b and Fig. 1d**).

To determine the stoichiometry of the GS-GN complex, we first performed mass photometry analysis of GS-GN and GS-GN(Y195F) mutant complexes, which enables mass measurements of single molecules in solution. Mass photometry measurements of the GS-GN(Y195F) complex showed a predominant species with an average molecular weight of 473 kDa, which is suggestive of a 4:4 stoichiometry (calculated mass of 485 kDa) (**Fig. 1f**). Analysis of the GS-GN(WT) sample identified a species with an average molecular weight of 534 kDa and the measured peak was broader than the non-glucosylated species (**Fig. 1f**). While mass photometry measurements lack the resolution to ascertain the precise molecular mass of heterogeneously glucosylated species, the observed increase in average molecular mass and overall distribution of the WT complex when compared to the Y195F complex is consistent with the observed higher molecular weight of WT GN1 glucosylated species (**Fig. 1d and Supplementary Fig. 3b**).

To understand how GS and GN interact and to reveal the overall shape of the GS-GN complex we performed negative stain electron microscopy (nsEM) of the WT and Y195F complexes. 2D class averages show two GN dimers, one on either side of a GS tetramer, for both WT and mutant complexes (**Fig. 1g**). Final 3D maps for both complexes are consistent with the 2D classes, and the reconstructed 3D EM density map can accommodate a GS tetramer flanked by two GN dimers (**Fig. 1h**). This nsEM confirms a 4:4 stoichiometry and is consistent with previous findings showing that GS can interact with four GN C-terminal peptides simultaneously^4,5,17^. Surprisingly, GN dimers do not engage the GS tetramer in an identical fashion, with one GN dimer tilted slightly towards GS and bringing it closer to one of the GS subunits (**Fig. 1h**). Collectively, these results provide the first glimpse of the glycogen initiating particle, where two GN dimers can engage a single GS tetramer.

### Phosphorylated human GS is in the inactive state

GS is regulated by both allosteric activation by G6P and inhibition via phosphorylation of its N- and C-terminal tails^2^ (**Fig. 1c**). Mechanistic and structural studies of yeast GS have elegantly dissected its allosteric activation by G6P^30,34^. However, GS structures to date were from protein preparations produced in bacterial expression systems and thus could not provide insights into the phospho-regulatory apparatus. Our GS-GN preparations are from eukaryotic expression systems and therefore provide an opportunity to study the inactive GS form. We confirmed that GS was phosphorylated at sites 2 (S8) and 3a (S641) and the enzyme preparation was inactive unless stimulated by G6P or dephosphorylation (**Fig. 2a and 2b**). Protein phosphatase 1 (PP1) and lambda protein phosphatase (lambda PP) treatment resulted in faster migration of GS by SDS-PAGE and also a reduction in signal detected by specific phosphorylation site antibodies (**Fig. 2a and Supplementary Fig. 4a**). Notably, we see only minor dephosphorylation of the GS-GN(Y195F) complex with PP1 alone, which was associated with a 5-fold increase in basal activity (-G6P) (**Fig. 2a and 2b**). We observed a 15-fold increase in basal activity when GS is dephosphorylated by both PP1 and lambda PP (**Fig. 2b**). The phosphorylated and dephosphorylated GS forms were similarly active after addition of G6P (**Fig. 2b**), which is consistent with studies using GS from endogenous sources^39,40^.

**Fig. 2.**
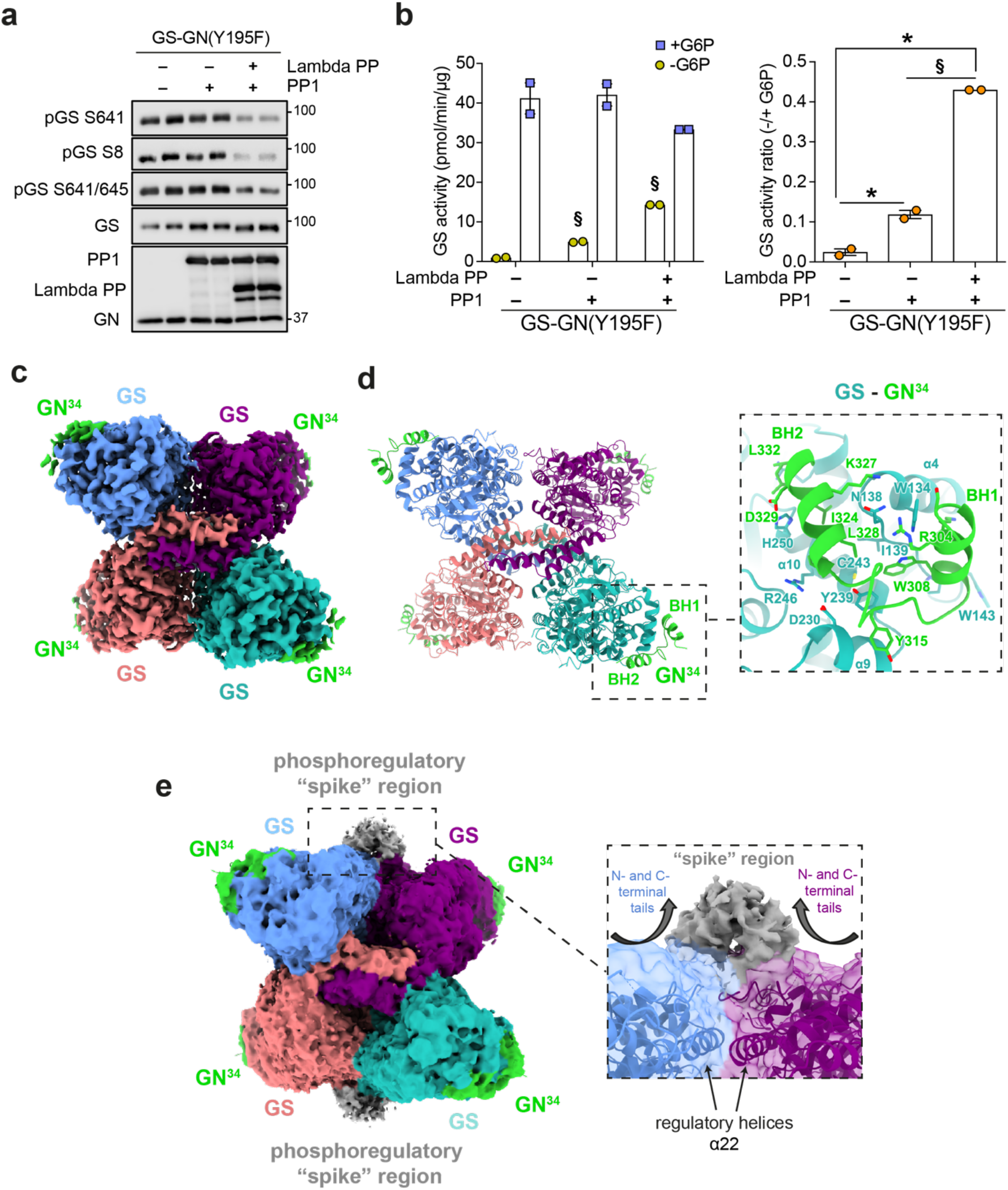
Cryo-EM structure of human GS-GN^34^ complex. **a** Immunoblot for the indicated human GS phosphorylation sites and total GS. **b** Activity of GS-GN(Y195F) with and without the addition of lambda protein phosphatase (lambda PP) and protein phosphatase 1 (PP1) (left) and -/+ G6P activity ratio (right). Upon G6P saturation, GS reaches similar activity levels regardless of phosphorylation state. Data are mean +/- S.E.M. from n=2 and representative of two independent experiments. One-way analysis of variance (Tukey’s post hoc test); §= *p*<0.05, +PP1 vs. +PP1+lambda PP (-G6P) (left). *= –PP1–lambda PP vs. +PP1 vs. +PP1+lambda PP, §= +PP1 vs. +PP1+lambda PP (right). **c** 2.6 Å cryo-EM map of the GS tetramer coloured by corresponding chain. Density corresponding to the GN^34^ C-terminal region is shown in green. **d** Human GS-GN^34^ cartoon model shown in ribbons coloured by corresponding chain (left). Interaction between GS and GN^34^ (right). **e** Unsharpened cryo-EM map shown at a lower threshold to visualise the “spike” region depicted in grey (left). The N- and C-terminal tails of two protomers converge and form the “spike” region (right).

To better understand the extent of phosphorylation we used tandem mass spectrometry (MS/MS) after proteolysis with either trypsin, chymotrypsin or elastase to map the phosphorylation sites of human GS. This resulted in a sequence coverage of 97%, which is higher than the 73%^35^ and 65%^36^ sequence coverage achieved in previous studies (**Table 1 and Supplementary Fig. 4b**). Our analysis identified canonical sites 2, 3a, 3b, 4 and 5 (S8, S641, S645, S653, S657), and also non-conventional sites (S412, S652, S727, S731). In addition, we could detect human GS site 2 (S8) phosphorylation by mass spectrometry for the first time in a recombinant enzyme preparation. Together, these results show that expression in insect cells is sufficient to achieve phosphorylation at multiple inhibitory sites and to provide suitable enzyme preparations to study inactive GS.

**Table 1:**
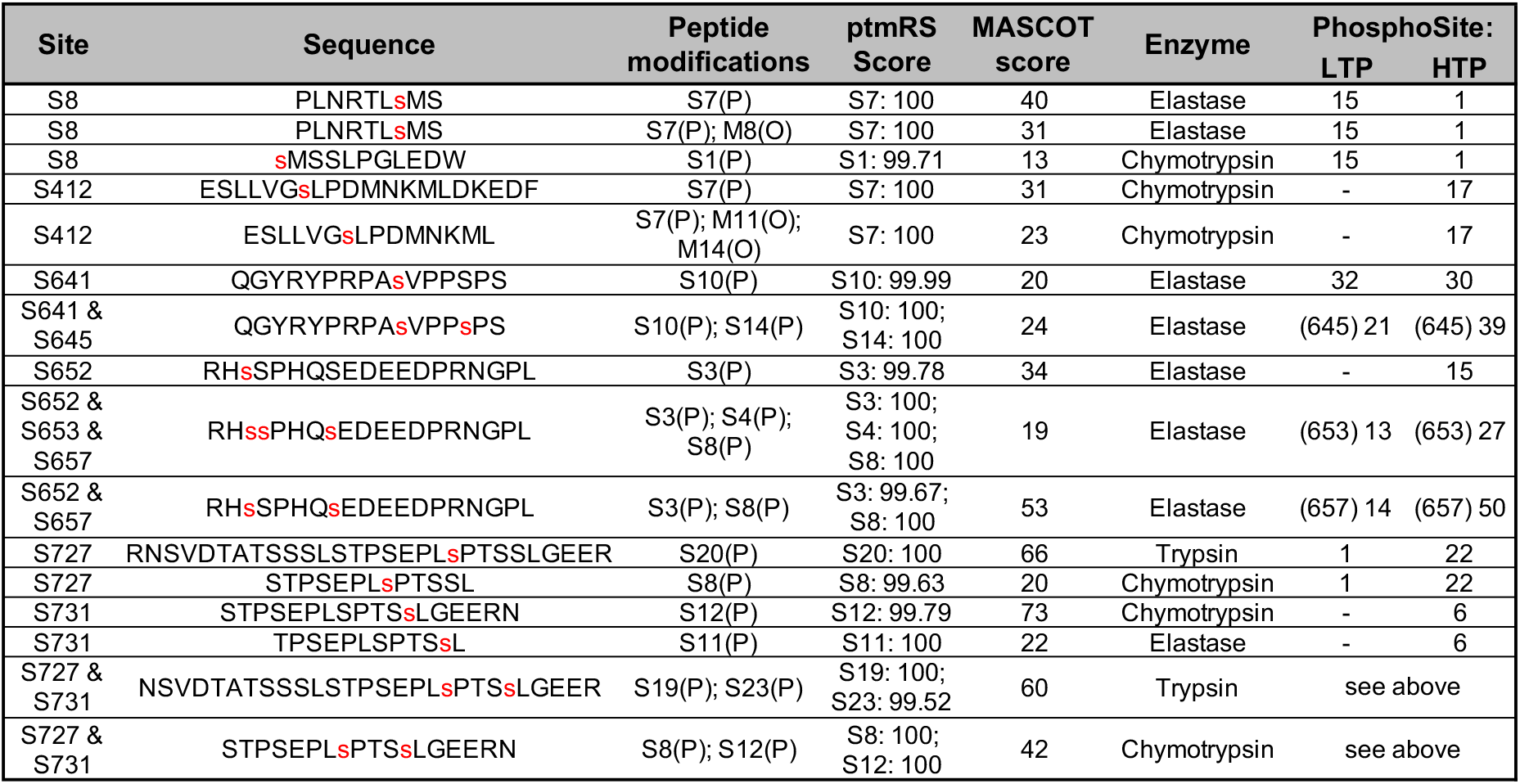
Summary of GS phosphorylation site analysis. Peptide modifications show either phosphorylation (P) or oxidation (O). PhosphoSite Plus (web-based bioinformatics resource) was used for comparison of our results with previous literature. LTP and HTP refer to low throughput site determination (methods other than mass spectrometry) and high throughput analysis (mass spectrometry only), respectively^41^.

### High resolution structure of human GS

Previous attempts to crystallise full-length GS in complex with full-length GN were unsuccessful^22^ leading us to pursue structural analysis using cryo-electron microscopy (cryo-EM). NsEM indicated that the position of each GN dimer is different suggesting flexibility of GN in the complex (**Fig. 1g and 1h**). Cryo-EM analysis of the GS-GN(Y195F) complex confirmed this GN flexibility as evidenced from the lack of GN signal in 2D class averages (**Supplementary Fig. 5a**) and subsequent 3D maps. Although we could detect the presence of GN after data processing without the application of symmetry averaging (**Supplementary Fig. 6c**), it was not possible to trace the connecting residues between the GN globular domain and the C-terminal GN^34^ region that binds GS. To gain a higher resolution structure for the human GS, we applied D2 symmetry and achieved a global resolution of 2.6 Å (EMDB-14587) (**Fig. 2c, Supplementary Fig. 5 and Supplementary Table 1**). The 3D reconstruction revealed a tetrameric arrangement of human GS in agreement with the crystal structures of the *C. elegans* GS and yeast GS enzymes, with root mean square deviation (RMSD) values of 1.1 Å (between 484 Cα atom pairs) and 0.9 Å (between 522 Cα atom pairs) respectively (**Fig. 2d, Supplementary Fig. 7a and 7b**). Structural analysis of the human GS-GN(WT) complex revealed a 6 Å map of the GS tetramer and comparing this to the GS structure from human GS-GN(Y195F) complex reveals no differences at this resolution (**Supplementary Fig. 6d and 6e**).

Density for the C-terminal GS interacting region of GN allows for model building of residues 300-332 (human GN^34^). Four GN peptides bind to the GS tetramer, and these residues form a helix-turn helix, where the first helix is denoted binding helix 1 (BH1) and the second as BH2 (**Fig. 2d**). This is consistent with the *C. elegans* GS-GN^34^ crystal structure^22^, with an RMSD value of 0.8 Å (between 30 Cα atom pairs) (**Supplementary Fig. 7c**). The interaction interface between human GS, namely α4, α9 and α10, and human GN^34^ is mediated by a combination of hydrophobic and hydrogen bonding interactions and is consistent with the interactions observed for GS-GN^34^ from *C. elegans*^22^ (**Fig. 2d and Supplementary Fig. 7c**).

### Mechanism of GS inactivation

A unique feature of metazoan GS is that both N- and C-terminal tails are phosphorylated, but the mechanism by which they participate in enzyme inactivation has remained elusive. We were able to build a model for the N-terminus starting from residue 13, and of the C-terminus up to residue 625, and then from 630-639 (chain A/C) and 630-642 (chain B/D), that could help understand the mechanisms of GS inactivation (PDB 7ZBN). The N- and C-terminal tails of each GS protomer lie almost parallel to each other, and travel side by side along the GS tetrameric core to reach the centre (**Fig. 3a**, right panels). Here, the C-terminal tail (chain A) meets the C-terminal tail from an adjacent GS protomer (chain B), which has travelled from the opposite direction (**Fig. 3a**, right panels). A 2.8 Å cryo-EM map of GS generated without the application of D2 symmetry averaging (EMDB-14587) (**Supplementary Fig. 6a and 6b**), suggests that one C-terminal tail disengages with the GS core earlier than the other C-terminal tail from the adjacent chain. The C-terminal tail from chain B continues to travel further across the regulatory helices than chain A, prior to traversing away from the core (**Fig. 3a**). This allows chain B to engage with the regulatory helices α22, specifically phosphorylated S641 interacting with residues R588 and R591, which come from two GS protomers to form a positively charged pocket we have termed the “arginine cradle” (**Fig. 3a and Supplementary Fig. 7d**). This is consistent with our phosphorylation mapping and immunoblotting data showing S641 is phosphorylated in our preparations (**Table 1, Supplementary Fig. 4b and Fig. 2a**).

**Fig. 3.**
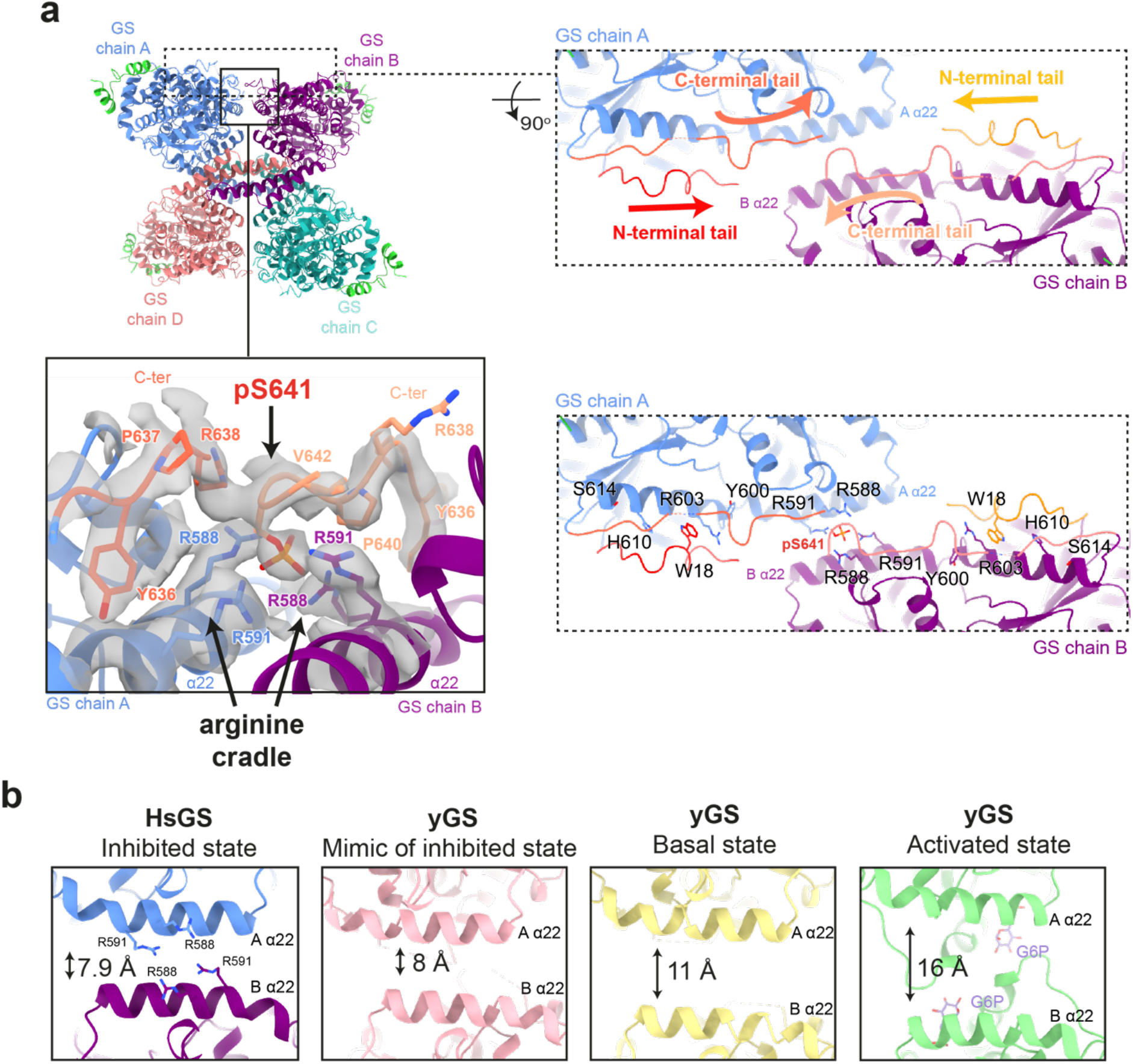
The phosphoregulatory region of human GS. **a** Human (Hs)GS-GN^34^ structure shown in ribbons (top left). The N- and C-terminal tails of one GS protomer (chain A) lie next to one another and move towards the adjacent protomer, meeting the N- and C-terminal tails from chain B. Arrows indicate continuation of cryo-EM density (top right). Electron density (C1 symmetry) for phosphorylated S641 (pS641) interacting with R588 and R591 on the regulatory helices α22 (bottom left). Residues that are interacting with the N- and C-terminal tails that are mutated in this study are shown (bottom right). **b** Comparison of distances between regulatory helices of adjacent monomers of HsGS (reported here), low activity inhibited mimic (PDB ID 5SUL), basal state (PDB ID 3NAZ) and G6P activated (PDB ID 5SUK) yeast crystal structure. Quoted distances were measured from Cα of Arg591 (chain A) and -Cα of Arg580 (chain B) of HsGS and corresponding yeast (yGS) residues.

S641 is a major phosphorylation site involved in the regulation of GS activity^40,42^, and interaction of pS641 with the arginine cradle in helix α22 shows the mechanism of inactivation of human GS through constraining the GS tetramer in a “tense state”. This interaction therefore provides a crucial activity switch mechanism from a tense (phosphorylated) state to a relaxed (G6P-bound) state^30^. The involvement of helices α22, which also interact with G6P via the nearby arginine residues R582 and R586^30^ (**Supplementary Fig. 8a**), provides a possible link between G6P-binding and its ability to override inactivation by phosphorylation.

The Rossmann fold domains of human GS were predicted to a high level of accuracy by AlphaFold^43^ (RMSD 1.0 Å between 575 Cα atoms), although the position of the N- and C-terminal tails does not agree entirely (**Supplementary Fig. 8c**). However, the position of S641 is consistent and overlays well with the phospho-S641 modelled in our cryo-EM structure (**Supplementary Fig. 8d**). This suggests that a Ser641 interaction with the arginine cradle may also be possible in the non-phosphorylated state, although the negative charge on the phosphate group would naturally provide stronger interactions with the positively charged arginine cradle.

### GS contains a dynamic “spike” region

Notably, the EM structures maps show density for an inter-subunit region that extends from the N- and C-termini of two adjacent GS protomers. These N- and C-termini contain the GS phosphoregulatory apparatus which meet and traverse away from the GS core (**Fig. 2e**). Analysis of this ~25 kDa region by focussed 3D classification (without applying symmetry) reveals that the region is highly flexible, as seen by the various different conformations (**Supplementary Fig. 5f and 5g**). Interestingly, these “spike” regions were present in all the refined classes, and suggests that GS exists as a continuum of structures with a core inactive tetramer and “dynamic spikes” buttressed on either side, thus preventing GS from adopting an open, active conformation.

To explore the flexibility and mobility of GS, we performed 3D variability analysis^44^ using cryoSPARC^45^. The dynamic movements of the “spike” region and concurrent movements of the GS tetramer are highlighted in Movie S1. Consistent with the focussed 3D classification, the “spike” is highly mobile, whereas only slight flexibility was observed within each GS protomer. This suggests a role of the “spike” region in constricting a tense state of the GS tetramer, and subsequently contributing to the GS regulation.

### Cross species comparison of GS structures

When comparing human GS to previous crystal structures of yeast GS, the distance between regulatory helices (α22) in adjacent monomers changes according to the activity state of GS (**Fig. 3b**). In the phosphorylated human GS structure, helices α22 lie 7.9 Å apart when measuring Cα-Cα distances from Arg591 on chain A and Arg580 on chain B (**Fig. 3b**). A similar measurement of the corresponding residues in the yeast proteins shows that helices α22 are furthest apart, at 16 Å, when G6P is bound and GS is in its high activity state, and this translates into better access for accepting the substrate^30,34^. When no G6P is bound and there is no phosphorylation, GS is in the basal state and the helices lie 11 Å apart^30^. In a yeast GS structure of a mimic of the inhibited state, where residues R589 and R592 were mutated to Ala and GS was produced in bacteria, the helices are closest together at 8 Å^34^. This is similar to the phospho-human GS, where phosphorylation appears to contribute to the closing of the regulatory helices constraining the GS tetramer and thus locking it in a tense, inactive state (**Fig. 3b**).

The position of the extreme N-terminus is noticeably different in human and *C. elegans* GS structures compared to yeast (**Supplementary Fig. 7e**). The majority of the first β-sheet in all structures is in a similar orientation, however human residues before 26 (residue 7 in yeast) move in the opposite direction to yeast (**Supplementary Fig. 7e**). This positioning of the human GS N-terminus is directed towards the regulatory helices α22. Previous structural investigation of *C. elegans* GS-GN^34^ suggested a hypothesis where phosphorylation could enable the N-terminus to engage with regulatory helices, as the N-terminus is also situated towards the regulatory helices^22^ (**Supplementary Fig. 7e**). Our structure of the human, phosphorylated enzyme supports this hypothesis, although the current density does not allow model building before residue 13. However, using LAFTER^46^ denoised maps to aid model building and electron density interpretation, some density for the N-terminus is present next to the regulatory helices, near to R579 and R580. This suggests that perhaps the N-terminal phosphorylation sites can also interact with the regulatory helices and/or nearby residues (**Fig. 3a and Supplementary Fig. 7f**).

Comparisons between human, *C. elegans* and yeast GS structures are consistent with the human structure in the inactive state. Each human GS protomer shows a closed conformation of its active site, and a regulatory loop, that only becomes ordered upon G6P binding, is disordered in the human structure (**Supplementary Fig. 8b and 8a**). Previous studies have suggested that phosphorylated tails may be able to engage the G6P binding site and directly compete with G6P. However, our EM density maps show no extra density within the G6P binding site (**Supplementary Fig. 8a**). Thus, we see no evidence to support the hypothesis that the phosphorylated tails interact with residues lining the G6P pocket to directly compete with G6P binding. Instead, we posit that the phosphoregulatory regions indirectly affect G6P binding by constraining the opening and closing of the GS tetramer. Collectively, our structural analyses support a model by which phosphorylated N- and C-terminal tails inhibit the GS tetramer by constraining a tense conformation through inter-subunit interactions.

### Dislodging the GS phosphoregulatory region

Due to the flexibility evident in the N- and C-terminal tails, we were unable to build phosphorylated residues in the cryo-EM map other than phospho-S641. However, we can see the beginning of the flexible phosphoregulatory “spike” region and residues from the GS “core tetramer” which interact with this regulatory region (**Fig. 3a**, bottom right panel). To investigate the relationship between allosteric regulation and inhibitory phosphorylation and elucidate the mechanism of inactivation, we mutated residues in GS that contact the beginning of the phosphoregulatory region. We selected residues which are not involved in G6P binding and mutated these in order to “dislodge” the regulatory tails (**Fig. 3a and Supplementary Fig. 8e**). If the phosphorylated tails are indeed holding GS in an inactivated state, weakening the interaction between the core tetramer and the N- and C-termini inhibitory regions should create an enzyme with higher basal activity in comparison to the WT. Consistent with our hypothesis, we observed a marginal increase in basal (-G6P) GS activity in R588A+R591A, Y600A, R603A, H610E and W18A mutants, that was reflective of the phosphorylated state at residues S8, S641 and S645 (**Fig. 4a and 4b**). These mutants were unaffected in terms of GN co-purification and with the exception of R588A+R591A mutant, they had similar melting (T_m_) profiles and oligomeric state to the WT GS complexes (**Supplementary Fig. 3e, Fig. 4c, Supplementary Fig. 9**). All mutants except Y600A could still be activated to similar levels to the WT upon addition of G6P (**Fig. 4a**).

**Fig. 4.**
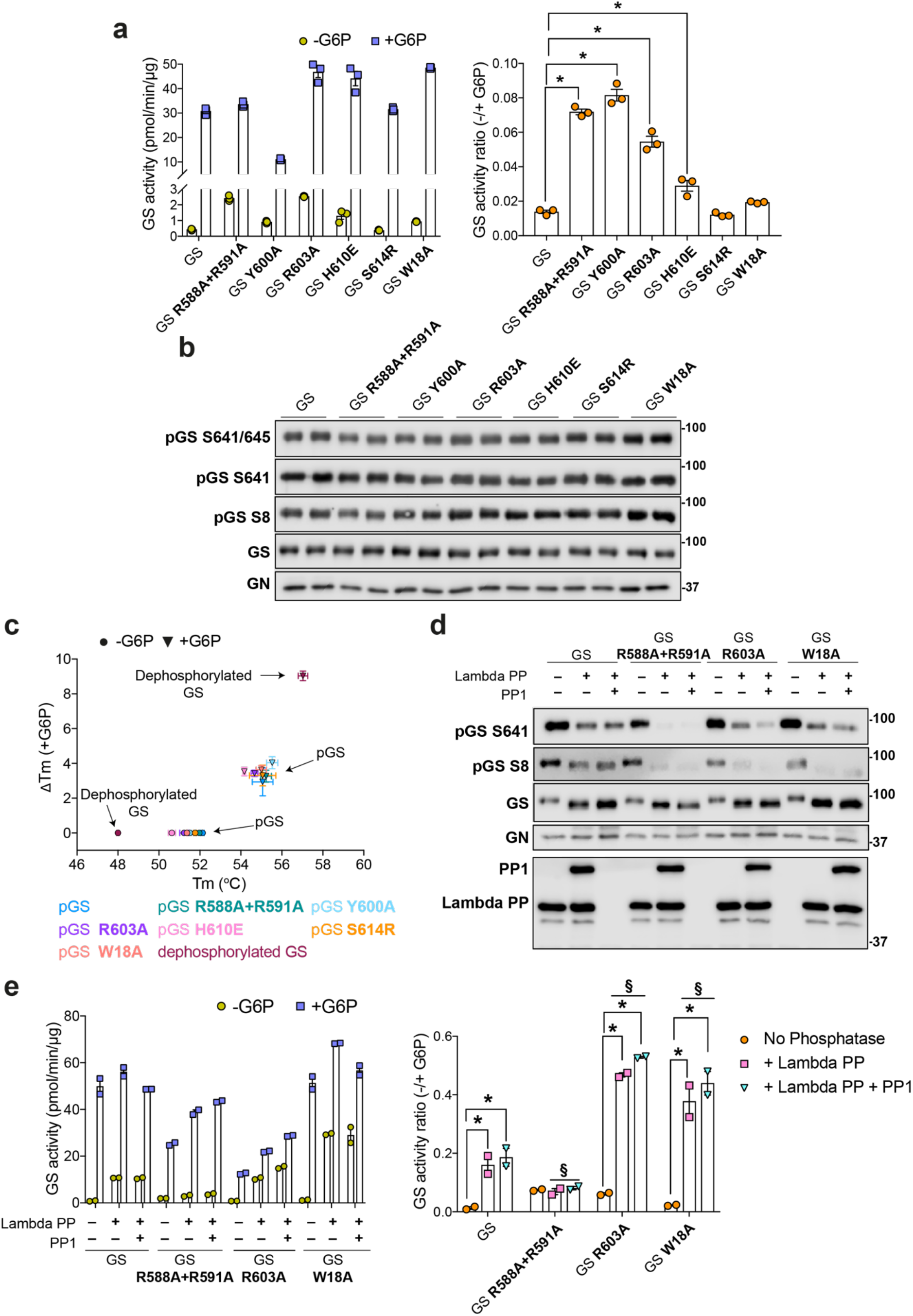
Dislodging the GS phosphoregulatory region increases basal activity and increases accessibility for phosphatases. **a** Activity of GS WT and indicated mutants in the GS-GN(Y195F) complex in the presence and absence of G6P (left) and -/+ G6P activity ratio (right). Data are mean +/- S.E.M from n=3 and representative of two independent experiments. One-way analysis of variance, (Tukey’s post hoc test); *= *p*<0.05 **b** Western blot for human GS phosphorylation sites S641/645, S641, S8, and total GS and GN. **c** Melting temperature (T_m_) of GS WT and mutants in the GS-GN(Y195F) complex. Changes in melting temperature upon addition of 12.5 mM G6P (ΔT_m_ = T_m_^+G6P^–T_m_^−G6P^). Data are mean +/- S.E.M from n=3 experiments carried out in technical duplicates (dephosphorylated GS) and triplicates (WT and mutant GS). **d** Western blots of GS WT and mutants in the GS-GN(Y195F) complex after dephosphorylation with PP1 and/or lambda protein phosphatase (lambda PP). **e** Activity of phosphorylated and dephosphorylated GS WT and indicated mutants (left) and -/+ G6P activity ratio (right). Data are mean +/- S.E.M from n=2 and representative of two independent experiments. Two-way analysis of variance (Tukey’s post hoc test); *p*<0.05 * = within groups, § = between groups: WT GS+lambda PP or +lambda PP+PP1 vs mutant GS +lambda PP or +lambda PP+PP1.

Upon addition of PP1 and lambda PP, the GS mutants R588A+R591A, R603A and W18A were more robustly dephosphorylated at S641 and S8 than WT GS (**Fig. 4d**), suggestive of increased exposure of the phospho-tails to phosphatases. For the W18A mutant, this dephosphorylation by both lambda-PP and PP1 resulted in over a 20-fold increase in basal activity, and also an approximately 3-fold increase in comparison to WT GS (**Fig. 4e**). The GS R603A-GN(Y195F) mutant has a basal activity similar to WT GS upon dephosphorylation. However, the robust dephosphorylation at S641 and S8 in GS R588A+R591A was not associated with an increase in activity (**Fig. 4e**). As described above, R588 and R591 lie on the regulatory helices and are also involved in inter-subunit interactions and form the arginine cradle that interacts with phospho-S641 (**Fig. 3a**). In addition, we noticed some dissociation of the GS R588A+R591A double mutant complex in mass photometry (**Supplementary Fig. 9c**). Therefore, the role of these residues in stabilising the GS tetramer may be the cause for the lack of rescue of activity upon dephosphorylation (**Fig. 4e**). Moreover, dephosphorylated GS had a markedly lower T_m_ (48 °C) than WT or mutant GS (**Fig. 4c**) supporting the idea that phosphorylation of the “spike” regions strengthens the inter-subunit interactions within the tetramer and holding the enzyme in the “tense” conformation.

## Discussion

For many decades human GS has remained elusive and resisted efforts for structural determination and characterization. Here, we provide structural and biochemical analysis of phosphorylated human GS in the full-length GS-GN complex. NsEM maps reveal two GN dimers binding to a GS tetramer, explaining the conformational plasticity of this octameric enzyme complex and the inner workings of how GS-GN cooperate to initiate glycogen synthesis (**Fig. 1 and Fig. 5**). The two GN dimers neighbouring a GS tetramer do not interact in an identical manner, with one GS dimer tilted closer towards GS in comparison to the other (**Fig. 1h**). This observed flexibility of GN may be aided by the variable length linkers connecting the catalytic domain and the C-terminal GN^34^ region that anchors GS. The precise functional relevance of this movement is yet to be explored, however the linker length was shown to govern glycogen particle size and molecular weight distribution *in vitro*^22^. Thus, the ability for GN to interact flexibly with GS may facilitate the wide range of size and distribution of glycogen particles seen in multiple species and tissues^4^.

**Fig. 5.**
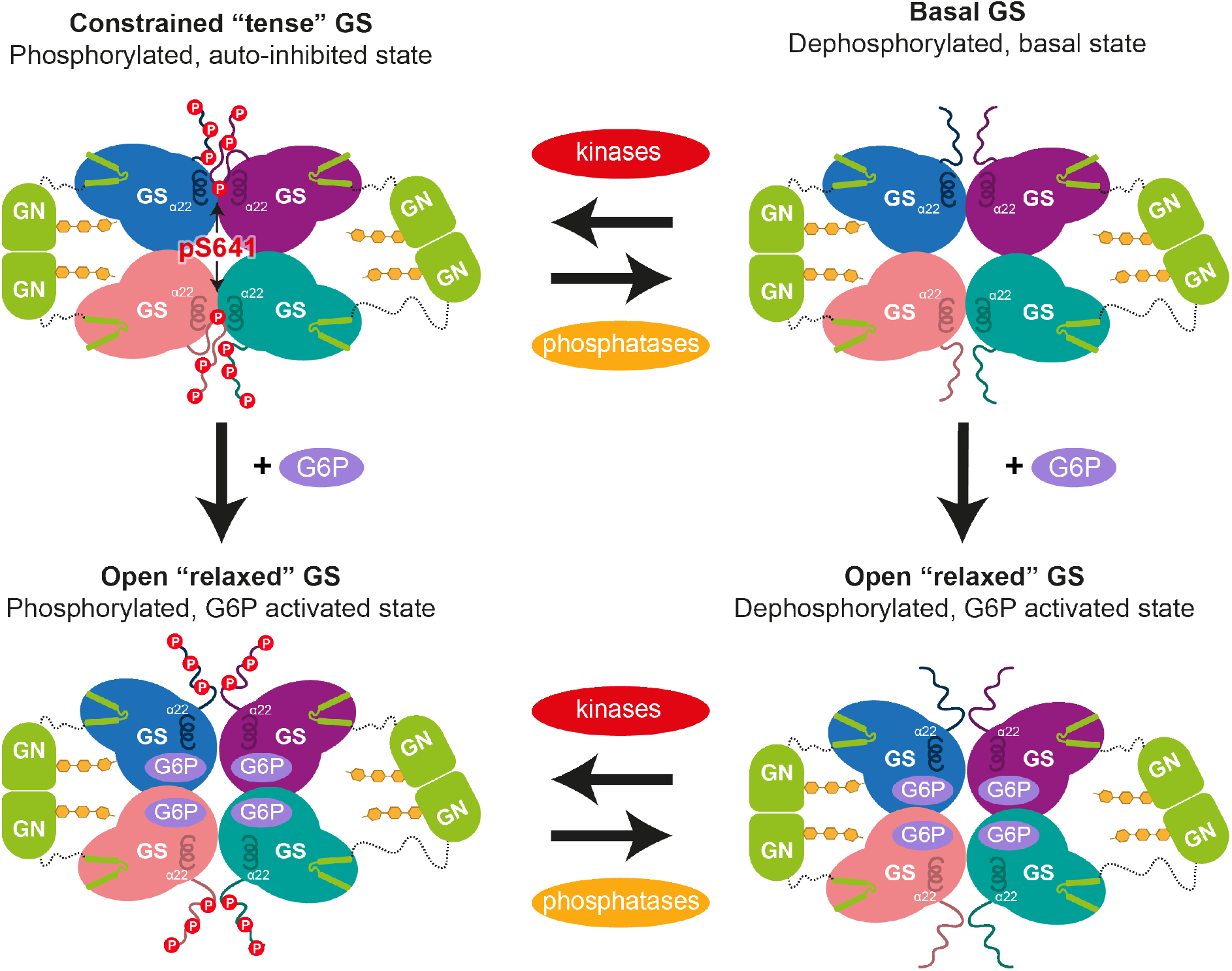
GS and GN cooperate to synthesize glycogen. Glucose is converted into glycogen through the action of glycogenin (GN), glycogen synthase (GS) and glycogen branching enzyme (GBE). GN interacts with GS to feed the initial glucose chain into the GS active site for elongation. GS is regulated by allosteric activation and inhibitory phosphorylation. Phospho-S641 (pS641) from one C-terminal tail interacts with the regulatory helices α22 to cause enzyme inhibition. This can be relieved by G6P, with or without phosphatases, to reach a high activity state. Kinases can phosphorylate GS to inhibit the enzyme.

Our human GS structure revealed phosphorylated S641 (site 3a) interacting with the regulatory helices α22. The electrostatic interactions between phospho-S641 and the arginine cradle could be strengthening the interaction between two GS protomers, thus constraining the GS tetramer and leading to an inactive enzyme (**Fig. 3a and 5**). Our structure also suggests a role for the N-terminal phosphorylation sites S8 and S11 (sites 2 and 2a) in the mechanism of human GS inactivation, as they presumably lie close to R579 and R580 on the regulatory helices (**Supplementary Fig. 7e and 7f**). Although model building before residue 13 was not possible, our analyses highlight an essential role for α22 in GS inhibition, specifically residues R579, R580, R588 and R591. This is consistent with previous studies showing that both N- and C-phosphorylation is required for inhibition of rabbit GS, as well as there being a significant role of site 3a and site 2/2a^23,42^. The role of the arginine cluster (residues 579-591, on the α22 helices) in GS regulation was investigated in yeast orthologues, revealing a role in G6P activation and suggesting a potential role in phosphorylation dependent inactivation^30,34,47,48^. Our inhibited human GS structure confirms that helix α22 is crucial for the phosphorylation dependent inactivation of GS, revealing that the same helix α22 is involved in both allosteric activation^30^ and covalent inhibition (**Fig. 3 and Fig. 5**).

GS phosphorylation sites lie outside of the catalytic core and within the N- and C-terminal tails (**Fig. 1c and Supplementary Fig. 2**). A comparison of human, *C. elegans* and yeast structures reveals considerable differences in the position of the tails (**Supplementary Fig. 7e**). In human GS, the C-terminus is responsible for most of the observable interactions with helix α22 via R588 and R591 residues, however, these residues are not conserved in *C. elegans* (**Supplementary Fig. 2**), perhaps explaining the positional differences between human and *C. elegans* GS tails. Interestingly, S641 (site 3a) is also not conserved in *C. elegans* GS, suggesting an evolutionary divergence, and hinting at additional mechanisms for *C. elegans* GS inactivation where the N-terminus interacts with helix α22. This potential exchange of interactions between N- and C-termini suggests a functional redundancy between the multiple phosphorylation sites.

The non-identical engagement of the C-terminal tails and the proximity of the N-terminus to the regulatory helices, as well as the flexibility of the “spike” region indicates coordination between the N- and C-termini of a single GS protomer, as well as between protomers (**Fig. 3a and Supplementary Fig. 7f and 5g**). Having one tail buttressed against the regulatory helix and the other steering away from the core may allow interchanging of the tails based on their level of phosphorylation, perhaps explaining why multiple phosphorylation sites are required. It may also aid/allow rapid dephosphorylation of GS, leading to an increase in GS activity and thus promoting glycogen synthesis (**Fig. 5**).

Elucidating the role of the inter-subunit domain that house the phosphorylation sites, through mutations that weaken the interactions between the core tetramer and the “spike” regions, resulted in basal activity equal to or higher than the WT, yet retaining activation by G6P (**Fig. 4a**). The GS Y600A mutant was not activated by G6P to the same extent as WT (**Fig. 4a**), and although Y600 does not directly bind to G6P, UDP or sugars^30,49,50^, it is possible that this hydrophobic residue is important for interdomain movements which are required for full GS activation.

Dephosphorylation of the GS-GN(Y195F) complex resulted in an increase in basal activity, yet there is little difference in the high activity (G6P-bound) state between phosphorylated and dephosphorylated complexes (**Fig. 2a and Fig. 4a**). This is in accordance with previous studies that demonstrate that G6P can overcome inhibition by phosphorylation and restore full activity^2^. C-terminal mutants R588A+R591A and R603A and the N-terminal W18A mutant were more easily dephosphorylated than WT (**Fig. 4d**) suggesting that dislodging of the phosphoregulatory region leads to phosphorylation sites being more accessible to phosphatases.

The robust dephosphorylation of GS R588A+R591A did not result in an increase in basal activity (**Fig. 4d and 4e**), which mirrors previous results in yeast^34^. The analogous mutations were used in yeast GS resulting in low basal activity, yet it could still be fully activated by G6P^34^. It was proposed that these residues in the regulatory helix are essential for keeping GS in a “spring loaded” intermediate state, and thus charge neutralization by mutation of arginine to alanine leads to the “tense” inactive state^34^. Our activity data agree with this as we don’t see an increase in basal activity despite dephosphorylation at S641 and S8, although we do see a marginal difference between this mutant and WT in the phosphorylated state (**Fig. 4a and 4e**). However, it is important to note that we also see some complex dissociation with the R588A+R591A mutant as evidenced by a larger dissociated complex peak in mass photometry (**Supplementary Fig. 9c**). It is possible that the dephosphorylated R588A+R591A mutant is also unstable and dissociates more easily than the phosphorylated mutant, resulting in a less active preparation.

Mutations of human GS1 and GS2 are common in glycogen storage diseases, and cluster within pockets of GS, affecting UDP-G, G6P and sugar binding. Some mutations affect the interaction between GS and GN^34^, consistent with the requirement of this interaction for glycogen synthesis^22^. The structure presented here will therefore provide a valuable resource to understand disease mutations. In addition, this new structure and increased understanding of GS regulation facilitates GS studies and its relevance in GSD, particularly Pompe and Lafora diseases where a reduction of glycogen levels could be beneficial^12–16^. The high resolution achieved here (2.6 Å) would undoubtedly be beneficial in efforts to design GS inhibitors that block G6P, substrate binding and/or GS-GN^34^ interaction.

GS has evolved a mechanism by which the phosphorylated N- and C-terminal “spike” regions hold GS in an inactive conformation that is relieved by dephosphorylation and/or G6P binding. We propose that the dynamic nature of these regulatory regions provides a functional redundancy mechanism and serves the purpose of exposing phosphorylated residues to phosphatases, thus allowing a “tuneable rheostat” instead of an on/off switch for regulating GS activity. Collectively, our analyses of the human GS-GN enzyme complexes reveal important mechanistic and structural details that could improve our understanding of GSDs.

## Materials and Methods

### Materials

Total GN antibody (S197C, third bleed) was obtained from MRC-PPU Reagents and Services. Total GS (#3898) and p-GS S641 (#47043) antibodies were from Cell Signaling Technologies. p-GS S641/S645 (#07-817) is from MerckMillipore. Affinity-purified p-GS S8 antibody (YZ5716) was custom-generated by (YenZym Antibodies Brisbane, CA, USA) by immunisation with a combination of phosphorylated peptides of the mouse GS1 (residues 2-14: PLSRSL-*S-VSSLPG-Ahx-C-amide, in which the prefix * denotes the phosphorylated residue) and human GS1 (residues 2-14: PLNRTL-*S-MSSLPG-Ahx-C-amide). Ahx and Cysteine (C) were added at the C terminal of the antigen peptides as linker/spacer and for conjugation to carrier protein, respectively. Antibody validation is shown in Supplementary Fig. 4c. Secondary antibodies (#711-035-152 and #713-035-147) were obtained from BioRad. Glucose-6-phosphate (G6P) (#10127647001) is from Roche. All other chemicals if not noted otherwise are from Sigma Aldrich.

### Cloning, protein expression and purification of GS-GN complex

Genes encoding human GS1 (HsGS:NM 002103) and human GN1 (HsGN:NM 001184720) mutant were cloned into pFL a vector^51^. A single 6 x His purification tag followed by a cleavable site was engineered at the N-terminus of GN WT or GN Y195F mutant. For co-expression of WT GS and mutants the genes encoding human GS1 and human GN1 (Y195F) were cloned in pFastBac vectors, both with a 6x His purification tag followed by a TEV site at the N-terminus. Recombinant bacmids were generated in DH10Bac™ cells. Virus amplification and protein expression, in *Spodoptera frugiperda* (Sf9) cells and *Trichoplusia ni* (Tni) cells respectively, were carried out using standard procedures^52^. For co-infection of pFastBac clones, a 10:1 ratio of the GS:GN P2 virus ratio was used. A PCR-based site directed mutagenesis was used to create the following mutants from the pFastBac GS1 construct: W18A, R588A+R591A, Y600A, R603A, H610E, S614R. All of the alterations were confirmed by DNA sequencing.

Cell pellets containing HsGS-GN, HsGS-GN(Y195F) and mutants were resuspended in lysis buffer (50 mM Tris-HCl pH 7.6, 300 mM NaCl, 20 mM imidazole, 10% glycerol, 0.075% β-mercaptoethanol, 1 mM benzamidine, 0.8 mM phenylmethyl sulfonyl fluoride (PMSF), 0.3 mg/mL lysozyme). Cells were lysed by sonication (1 second on, 3 seconds off for a total of 5 minutes) on ice and the lysate was cleared by centrifugation at 35,000*g* for 30 minutes at 4 °C. The clarified lysate was sonicated again (1 second on, 3 seconds off for a total of 1 minute), followed by filtering with a 0.45 μM filter (MerckMillipore). Filtered lysate was loaded onto a pre-equilibrated 1 mL or 5 mL HisTrap HP column (GE Healthcare) charged with Ni^2+^. The loaded column was washed with four column volumes (CV) of low salt buffer (50 mM Tris-HCl pH 7.6, 300 mM NaCl, 20 mM imidazole, 10% glycerol, 0.075% β-mercaptoethanol, 1mM benzamidine), followed by four CV washes of high salt buffer (50 mM Tris-HCl pH 7.6, 500 mM NaCl, 20 mM imidazole, 10% glycerol, 0.075% β-mercaptoethanol, 1 mM benzamidine) and finally 4 CV washes in low salt buffer. The column was then attached to the AKTA system (GE Healthcare) and washed with low salt buffer. The protein was then eluted by applying an imidazole gradient with elution buffer (50 mM Tris-HCl pH 7.6, 300 mM NaCl, 300 mM imidazole, 10% glycerol, 0.075% β-mercaptoethanol, 1 mM benzamidine). The fractions containing protein were analysed by SDS-PAGE and then pooled and dialysed overnight (10,000 MWCO SnakeSkin dialysis tubing (Thermo Scientific)) at 4 °C in dialysis buffer with TEV protease added (50 mM Tris-HCl pH 7.6, 150 mM NaCl, 20 mM imidazole, 10% glycerol, 0.075% β-mercaptoethanol, 1 mM benzamidine). The dialysed protein was re-loaded onto the HisTrap column equilibrated with low salt buffer, for a Ni subtraction step. TEV cleaved protein was eluted in the flow through and low salt washes. The flow through and first low salt wash were pooled and concentrated using a VIVASPIN20 30,000 MWCO (Sartorius, Generon), followed by centrifugation at 17,000*g* for 15 minutes at 4 °C. Protein was then injected onto a 16/600 or 10/300 Superdex200 column (GE Healthcare) equilibrated with gel filtration buffer (25 mM HEPES pH 7.5, 150 mM NaCl, 1 mM TCEP, 10% glycerol). Fractions containing protein were analysed by SDS-PAGE, and fractions containing GN were pooled, concentrated and stored at −80 °C. Some fractions containing GS-GN complex were stored separately at −80 °C and the remaining protein was pooled and concentrated before being stored at −80 °C. Proteins were visualized by Coomassie blue staining, and glucosylated species were detected using the periodic acid-Schiff (PAS) method (Glycoprotein staining kit, Thermo Scientific).

### *In vitro* dephosphorylation of GS-GN

Protein phosphatase 1 (PP1) and lambda protein phosphatase (lambda PP) were bought from MRC PPU Reagents and Services. Both have an N-terminal GST tag and lambda PP also has a C-terminal 6x His tag. GS-GN complex was dephosphorylated in reactions containing equal amounts of PP1 and lambda PP in 25 mM HEPES pH 7.5, 150 mM NaCl, 1 mM TCEP, 1 mM MnCl_2_ and 10% glycerol for 30 minutes at 30 °C. For subsequent differential scanning fluorimetry experiments, the phosphatases were removed by incubating the reactions with GST beads for 1 hour at 4 °C. Reactions were then passed through an equilibrated 0.45 μm Spin-X column (Costar, 0.45 μm cellulose acetate) and eluted by centrifugation at 16,000*g* for 2 minutes.

### *In vitro* deglycosylation

GS-GN was incubated with α-amylase from human saliva (Sigma) to deglycosylate the complex. Reactions contained 4 μM GS-GN with either 500 mU or 1 U α-amylase in buffer containing 50 mM HEPES pH 7.5, 150 mM NaCl, 1 mM TCEP and 5 mM CaCl_2_. Reactions were incubated for 30 min, 1 hour or 2 hours at 37 °C and terminated by the addition of SDS-PAGE loading dye.

### Negative stain electron microscopy – grid preparation and data collection

HsGS-GN WT and Y195F were diluted in buffer (25 mM HEPES pH 7.5, 150 mM NaCl, 1 mM TCEP, 10% glycerol) to concentrations between 0.01 and 0.02 mg/mL immediately before grid preparation. Carbon-coated copper grids (Formvar/Carbon 300 mesh Cu, Agar Scientific) were glow-discharged for 30 seconds, 10 mA and 0.39 mBar pressure (PELCO easiGlow, Ted Pella). Grids were incubated for 1 minute with 6 μL sample, washed with H_2_O three times and stained twice with 2% w/v uranyl acetate for 20 seconds. Excess liquid was removed by blotting with filter paper. Data was collected on a FEI Technai F20 electron microscope operated at 120 keV, equipped with a FEI Ceta (CMOS CCD) camera.

### Negative stain electron microscopy – data processing

RELION 3.0 was used for processing of negative stain-EM data^53^. Real-time contrast transfer function (CTF) parameters were determined using gCTF^54^. Approximately 2,000 particles were manually picked, extracted with a box size of 104 Å^2^, then subjected to reference-free 2D classification to produce initial references to be used for auto-picking. The parameters for auto-picking were optimized and 92,580 particles were extracted. The extracted particles were used for iterative rounds of reference-free 2D classification. Based on visual inspection, best quality 2D average classes were selected to generate a *de novo* 3D initial model, which was used as a reference in unsupervised 3D classification. These classes were then subjected to 3D refinement to generate a final EM density map.

### Cryo-electron microscopy – grid preparation and data collection

Quantifoil R2/2 Cu300 or Quantifoil R1.2/1.3 Cu300 (Quantifoil Micro Tools) grids were glow-discharged using the GloQube plasma cleaner (Quorum) at 40 mA for 30 seconds, for GS-GN(Y195F) and GS-GN respectively. A FEI Vitrobot IV was equilibrated at 4 °C at 100% relative humidity. GS-GN(Y195F) complex was diluted in buffer (25 mM HEPES pH7.5, 150 mM NaCl, 1 mM TCEP) to 0.8 mg/mL (6.59 μM) containing 1.5% glycerol immediately before 3 μL was added to the grid. GS-GN was diluted to 0.36 mg/mL (2.97 μM) containing 8% glycerol. This was followed by immediate blotting and plunge-freezing into liquid ethane cooled by liquid nitrogen.

All data was collected on a FEI Titan KRIOS transmission electron microscope at 300 keV. For GS-GN(Y195F), A FEI Falcon IV direct electron detector with an energy filter (10 eV) was used in counting mode^55^. A dose per physical pixel/s used resulting in a total dose of 34.8 e/Å^2^, fractionated across 128 EPU frames. This was then grouped into 21 frames, resulting in a dose per frame of 0.8 e/Å^2^. Magnification was 165,000x resulting in a pixel size of 0.71 Å/pixel. Eight exposures per hole was taken and the defocus values ranged from −1 μm to −2.2 μm. 20,841 movies were recorded using the EPU automated acquisition software (v2.13).

For GS-GN, a FEI Falcon III direct electron detector was used in integrating mode^55^. The total electron dose was 85 e/Å^2^, a magnification of 75,000x was used and a final calibrated object sampling of 1.065 Å/pixel. Each movie had a total exposure time of 1.6 seconds, collected over 47 fractions with an electron dose of 1.8 e/Å^2^ per fraction. One exposure per hole was taken and the defocus values ranged from −1.7 μm to −3.1 μm. 3,009 movies were recorded using the EPU automated acquisition

### Cryo-electron microscopy – data processing

For GS-GN(Y195F), drift-corrected averages of each movie were created using MotionCor2^56^ and real-time contrast transfer function (CTF) parameters were determined using CTFFIND-4.1^57^. Both motion correction and CTF estimation were carried out on-the fly^55^. 1,883,188 particles were picked using the PhosaurusNet general model in crYOLO^58^ v1.6.1. Particles were imported into RELION 3.1 and extracted and binned by 2. These particles were subjected to 2D classification. 1,188,332 particles selected after 2D classification were subjected to 3D classification, applying D2 symmetry. Carrying all “good”/unambiguous classes forward, 739,232 particles were un-binned to a box size of 288 pixels and subjected to 3Drefinement and postprocessing, generating a map at 2.92 Å. Followed by iterative rounds of per particle contrast transfer function refinement and Bayesian particle polishing to generate a map at 2.62 Å (**Supplementary Fig. 5**). Final resolutions were determined using the gold-standard Fourier shell correlation criterion (FSC=0.143). Local resolution was estimated using the local resolution feature in RELION.

To prevent interpretation of any artefacts created by applying D2 symmetry, the data was also processed in C1 symmetry (**Supplementary Fig. 6**). The same particles after 2D classification were subjected to 3D classification without applying symmetry. High quality classes containing 783,177 particles were un-binned to a box size of 288 pixels and then subjected to 3D refinement and postprocessing, to generate a 3.1 Å map. Following iterative rounds of contrast transfer function refinement and Bayesian particle polishing to generate a 2.8 Å map.

To elucidate the movement of phosphoregulatory regions, an alignment free 3D classification with a mask containing the “spike” density was performed, using a regularisation parameter T of 60^59^ (**Supplementary Fig. 5f and 5g**).

To explore the heterogeneity in the dataset, the 3D variability analysis^44^ tool in cryoSPARC v3.2.0^45^ was used. The 739,232 particles after 3D classification were imported into cryoSPARC and homogenous refinement with a mask with no symmetry application was performed. The subsequent particles were used in the 3D variability analysis to solve 3 modes, using C1 symmetry and with a filter resolution of 5 Å. The subsequent frames were visualized in Chimera^60^. 1 mode is shown in **Movie S1**.

For GS-GN, drift-corrected averages of each movie were created using MotionCor2^56^ and real-time contrast transfer function (CTF) parameters were determined using gCTF^54^. 250,250 particles were picked using the PhosaurusNet general model in crYOLO^58^ v1.3.5. Particles were then imported into RELION 3.0^53^, extracted with a box size of 220 pixels and subjected to reference-free 2D classification. 84,557 particles selected after 2D classification were subjected to 3D classification. 36,972 particles (from 2 classes) were subjected to 3D refinement and postprocessing, generating a map at 6.0 Å (**Supplementary Fig. 6e and 6e**).

### Model building and refinement

A preliminary model of human GS was generated by AlphaFold^43^ (accessed 1 October 2021) and a preliminary model of last 34 residues of human GN was created by Phyre2^61^. These preliminary models were rigid body fitted into the cryo-EM density in UCSF Chimera^60^. The model was then built using iterative rounds of manual building in COOT^62^ and real space refinement in PHENIX v1.19^63^.

### Visualisation, structure analysis and sequence alignments

Visualisation and structure analysis were performed using ChimeraX^64^ or Chimera^60^. Multiple sequence alignments were performed using MUSCLE^65^ and displayed and edited using ALINE v1.0.025^66^.

### Mass photometry

Mass photometry experiments were performed using a Refyn One^MP^ mass photometer. Immediately prior to mass photometry measurements, proteins were diluted in 25 mM HEPES pH 7.5, 150 mM NaCl, 1 mM TCEP for a final concentration of 50 nM. For each measurement, (16 μL) buffer was added to a well and the focus point was found and adjusted when necessary. Protein (4 μL) was then added to the buffer droplet, the sample was mixed and movies of 60 seconds were recorded using AcquireMP. Data were analysed using DiscoverMP.

### Differential Scanning Fluorimetry

Thermal shift assays were performed using an Applied Biosystems QuantStudio 3 Real-Time PCR system. SYPRO™ Orange (Invitrogen) was used as a fluorescence probe. Proteins were diluted in 25 mM HEPES pH 7.5, 150 mM NaCl, 1 mM TCEP to a final concentration of 1 μM. Varied concentrations of G6P were added and the reaction was incubated at room temperature for 30 minutes. SYPRO Orange was diluted in 25 mM HEPES pH 7.5, 150 mM NaCl, 1 mM TCEP to a final concentration of 2.5 X, in a total reaction volume of 20 μL. The temperature was raised in 0.018 °C intervals from 20 °C to 95 °C. Data were analysed using Protein Thermal Shift™ v1.4.

### Tandem mass spectrometry

Concentrated purified protein complexes (6.75 μg) were diluted 30-fold in 25 mM ammonium bicarbonate pH 8.0 before being subject to reduction with dithiothreitol and alkylation with iodoacetamide, as previously described^67^. The eluent was equally divided into three for digestion with either: 33:1 (w/w) trypsin gold (Promega), 25:1 (w/w) chymotrypsin (Promega), or 10:1 (w/w) elastase (Promega), using the manufacturer’s recommended temperatures for 18 hours with 600 rpm shaking. Digests were then subject to in-house packed, strong cation exchange stage tip clean-up, as previously described by^68^. Dried peptides were solubilized in 20 μl of 3% (v/v) acetonitrile and 0.1% (v/v) TFA in water, sonicated for 10 minutes, and centrifuged at 13,000*g* for 15 minutes at 4 °C being separated using an Ultimate 3000 nano system (Dionex) by reversed-phase HPLC, over a 60-minute gradient, as described in^67^. All data acquisition was performed using a Thermo Orbitrap Fusion Lumos Tribrid mass spectrometer (Thermo Scientific), with higher-energy C-trap dissociation (HCD) fragmentation set at 32% normalized collision energy for 2+ to 5+ charge states. MS1 spectra were acquired in the Orbitrap (60K resolution at 200 *m/z*) over a range of 350 to 1400 *m/z*, AGC target = standard, maximum injection time = auto, with an intensity threshold for fragmentation of 2e^4^. MS2 spectra were acquired in the Orbitrap (30K resolution at 200 *m/z*), AGC target = standard, maximum injection time = dynamic. A dynamic exclusion window of 20 seconds was applied at a 10 ppm mass tolerance. Data was analysed by Proteome Discoverer 1.4 using the UniProt Human reviewed database (updated April 2020) with fixed modification = carbamidomethylation (C), variable modifications = oxidation (M) and phospho (S/T/Y), instrument type = electrospray ionization–Fourier-transform ion cyclotron resonance (ESI-FTICR), MS1 mass tolerance = 10 ppm, MS2 mass tolerance = 0.01 Da, and the *ptmRS* node on; set to a score > 99.0.

### Protein Identification Mass Spectrometry

10 μg of purified protein was separated by SDS-PAGE (10% resolving, 4% stacking) before colloidal Coomassie staining overnight and thorough washing in milliQ water^**69**^. A scalpel was used to excise the major band at ~85 kDa, and incremental bands spanning 43-55, 55-72, 95-130 and 130+ kDa. Bands were washed in 500 μL HPLC H_2_O for 10 minutes shaking at 1500 rpm, room temperature. Bands were then washed in 500 μL of 100 mM ammonium bicarbonate with water bath sonication, as before, for 10 minutes, before an equal volume of HPLC acetonitrile was added and sonication repeated. Previous two wash steps were repeated until the gel pieces were clear. 100 μL of reduction solution (4 mM dithiothreitol in 50 mM ammonium bicarbonate) was added to each gel slice and heated to 60 °C for 10 minutes with 600 rpm shaking. A final concentration of 16.4 mM iodoacetamide was added and incubated in darkness at room temperature for 30 minutes, before quenching by addition of 100 mM dithiothreitol to make a final concentration of 7 mM. All liquid was removed before dehydrating the gel slice by addition of 100 μL HPLC acetonitrile and shaking 1500 rpm at room temperature for 15 minutes. Dehydrydration was repeated until gel slices were opaque white and left open lid to dry completely (~15 minutes). 0.5 μg of trypsin in 40 mM ammonium bicarbonate was added to the dehydrated gel slices and incubated room temperature for 15 minutes. Residual liquid was removed and 100 μL of incubation solution (40 mM ammonium bicarbonate, 5% acetonitrile) added and incubated overnight at 37 °C with 600 rpm shaking. An equal volume of acetonitrile was added and left to shake for an additional 30 minutes. Gel slices were briefly centrifuged and supernatant collected. Supernatant was dried to completion, resuspended and analysed by LC-MS/MS as described before.

### Glycogen synthase activity assay

1 μg of purified protein was diluted in ice cold lysis buffer (270 mM sucrose, 50 mM Tris-HCl (pH 7.5), 1 mM EDTA, 1 mM EGTA, 1% (v/v) Triton X-100, 20 mM glycerol-2-phosphate, 50 mM NaF, 5 mM Na_4_P_2_O_7_, 1 mM DTT, 0.1 mM PMSF, 1 mM benzamidine, 1 mg/mL microcystin-LR, 2 mg/mL leupeptin, and 2 mg/mL pepstatin A) to a total volume of 100 μL. 20 μL of the protein was added to 80 μL of the assay buffer (25 mM Tris-HCl (pH 7.8), 50 mM NaF, 5 mM EDTA, 10 mg/ml glycogen, 1.5 mM UDP-glucose, 0.125% (v/v) β-mercaptoethanol and 0.15 mCi/mmol D-[^14^C]-UDP-glucose (American Radiolabelled Chemicals, Inc., ARC 0154) with 0 and 12.5 mM G6P. Reactions were incubated for 20 minutes at 30 °C with mild agitation at 300 rpm. The reactions were stopped by spotting 90 μL of the reaction mix onto 2.5 cm x 2.5 cm squares of filter paper (Whatman 3MM) which were immediately immersed in ice cold 66% ethanol and left to incubate with mild agitation for 20 minutes. The filter papers were washed thrice more in 66% ethanol for 20 minutes each and rinsed in acetone. The dried filters were subjected to scintillation counting.

### Statistical Analysis

Data are reported as mean ± standard error of the mean (SEM) and statistical analysis was performed using GraphPad Prism software. As indicated in the respective figure legends, one-way or two-way analysis of variance was performed with Tukey’s post hoc test. Statistical significance was set at *p*<0.05.

### Immunoblotting

Purified proteins were denatured in Laemmli buffer at 100 °C for 5 minutes. 100 ng of the protein was separated by SDS-PAGE on 4-10% gel and transferred onto nitrocellulose membranes (#926-31090, LiCOR). Membranes were blocked for 45 minutes at room temperature in 5% skim milk (Sigma, 1153630500) followed by incubation in TBST (10 mM Tris (pH 7.6), 137 mM NaCl, and 0.1% (v/v) Tween-20) containing 5% (w/v) BSA and the primary antibody overnight at 4 °C. The membranes were incubated for 45 minutes in HRP conjugated secondary antibodies diluted 1:10,000 in 3% skim milk, after extensive washing in TBST. The membranes were imaged using enhanced chemiluminescence (ECL) reagent (GE Healthcare). For total protein staining of blots, Revert™700 Total Protein Stain (LiCOR) was used.

For the validation of pGS S8 antibody (YenZyme, 1^st^ cycle, YZ7516) HEK293FT cells were co-transfected with 900 ng of GS (WT) or GS S8A along with GST tagged-GN. After 40 hours, transfected cells were harvested for protein. The blots were probed with pGS S8 antibody incubated with 20 μg of GS peptide (**Supplementary Fig. 4c**).

## Data availability

The cryo-EM maps have been deposited in the Electron Microscopy Data Bank under the accession code EMDB-14587. Coordinates have been deposited in the Protein Data Bank under the accession code 7ZBN.

## Author contributions

L.M. performed molecular biology, protein production, electron microscopy, differential scanning fluorimetry and mass photometry experiments, D.B. performed glycogen synthase activity and western blot assays, L.A.D. performed phosphorylation mapping and mass spectrometry experiments and C.B., S.V. and D.P.M. provided support with structural biology and protein production. L.M. and E.Z. drafted the manuscript with input from D.B., K.S., L.A.D. and C.E.E. and all authors revised it. J.P., C.E.E., C.H., J.B., N.A.R., H.K., K.S. and E.Z. provided supervision and project management. E.Z., K.S., C.E.E., C.H. and J.B. designed and interpreted data in consultation with all authors.

## Acknowledgments

L.M. is supported by an MRC Discovery Medicine North (DiMeN) iCASE studentship co-funded by UKRI and Vertex Pharmaceuticals. D.B. is supported by an International Postdoctoral Fellowship funded by the Novo Nordisk Foundation (NNF) Center for Basic Metabolic Research. C.E.E. and L.A.D. were supported by grants from BBSRC (BB/S018514/1, BB/M012557/1, BB/R000182/1). E.Z. was supported by a Sir Henry Dale Fellowship from the Wellcome Trust & Royal Society (200523/Z/16/Z) and Royal Society grant RG170407. The Astbury cryo-EM Facility is funded by a University of Leeds ABSL award and Wellcome Trust grants (108466/Z/15/Z and 221524/Z/20/Z). The NNF Center for Basic Metabolic Research is an independent Research Center at the University of Copenhagen, Denmark, and partially funded by an unconditional donation from NNF (Grant number NNF18CC0034900).

