## Supplementary Information for "Mechanism of glycogen synthase inactivation and interaction with glycogenin"

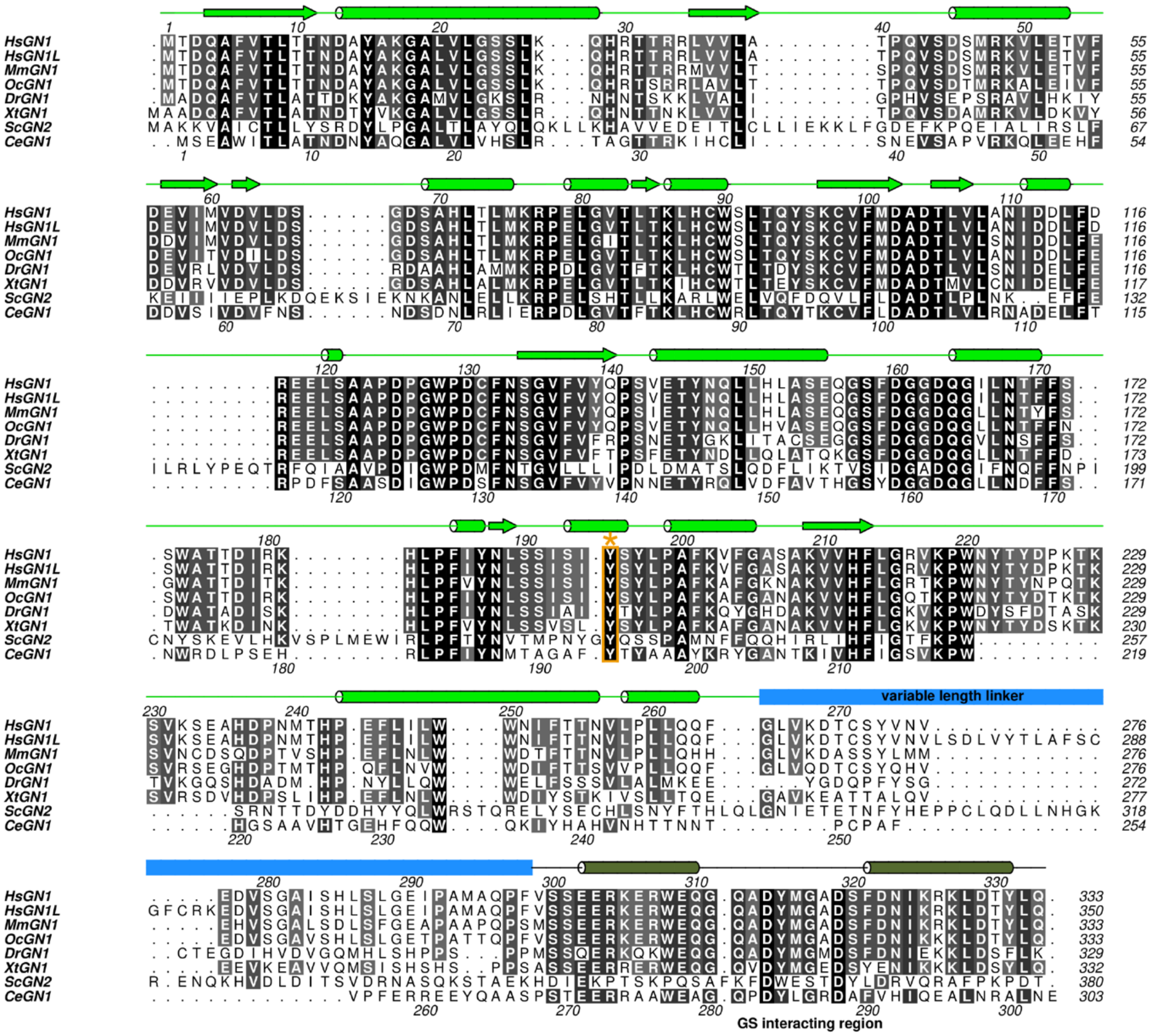

**Supplementary Fig. 1 Sequence conservation of glycogenin**

Multiple sequence alignment of GN from the indicated species (black = conserved, white = not conserved). Sequence alignment was performed with MUSCLE and edited and displayed using ALINE. The predicted secondary structure for HsGN is shown at the top of the alignment and coloured in green (analysed by PSIPRED). The human and *C. elegans* GN sequences are numbered at the top and bottom of the alignment respectively. The variable length linker is coloured in blue, the C-terminal GS interaction region is coloured in dark green and the tyrosine that is auto-glucosylated (Y195 in human) is boxed in orange and labelled with an asterix. *Hs, H. sapiens; Sc, S. cerevisiae; Ce C. elegans; Mm, M. musculus; Xt, X. tropicalis; Oc, O. cuniculus; Dr, D. rerio.*

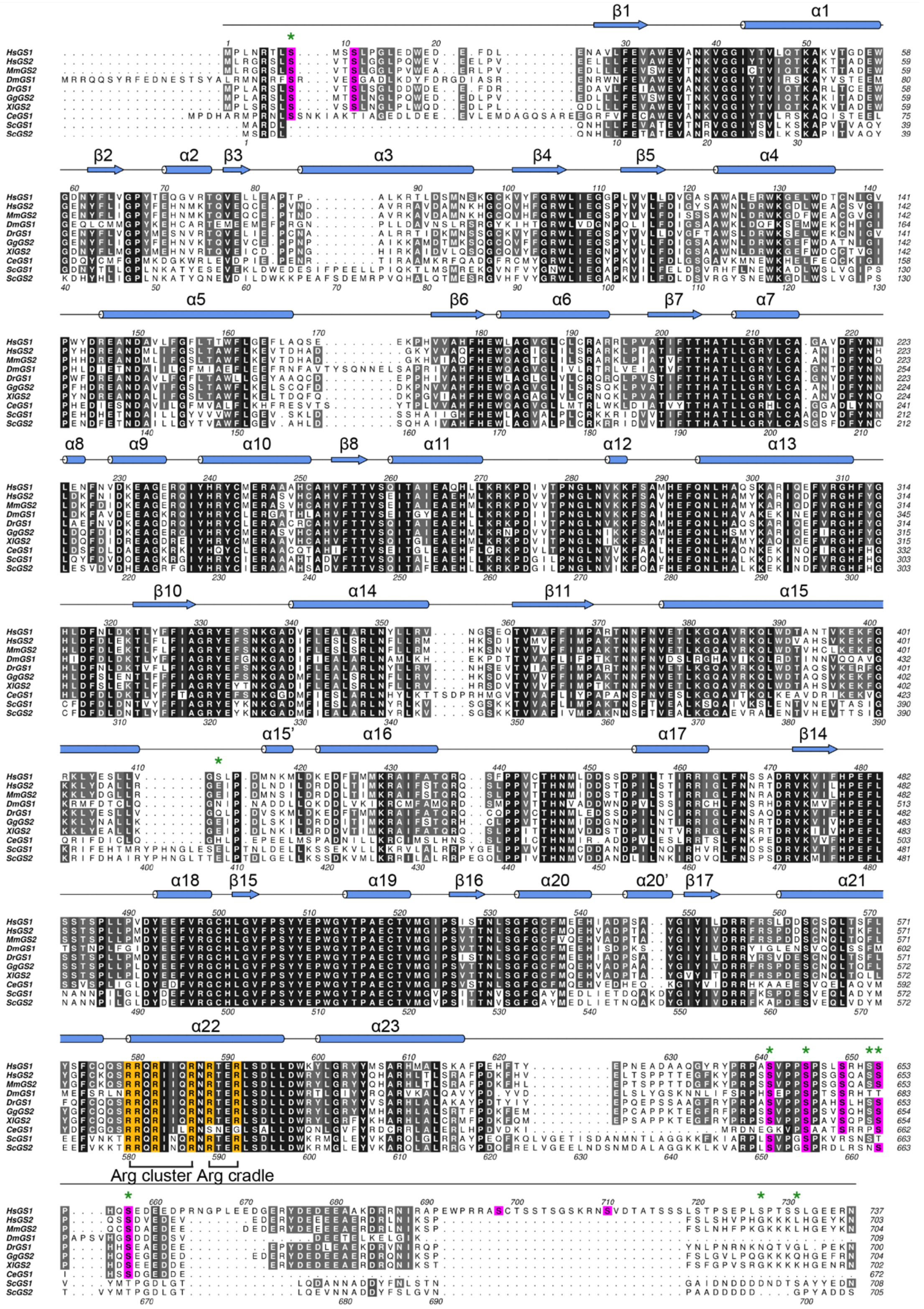

**Supplementary Fig. 2 Sequence conservation of glycogen synthase**

Multiple sequence alignment of GS from the indicated species (black = conserved, white = not conserved). Sequence alignment was performed with MUSCLE and edited and displayed using ALINE. The secondary structure for HsGS is shown at the top of the alignment and coloured in blue. The human and yeast GS sequences are numbered at the top and bottom of the alignment respectively. Known *in vivo* phosphorylation sites are shown in magenta. Residues that form the arginine cluster and arginine cradle are coloured in orange and labelled, and the phosphorylated residues identified by phosphomapping experiments in this study are shown by green stars. *Hs, H. sapiens; Sc, S. cerevisiae; Ce C. elegans; Dm, D. melanogaster; Mm, M. musculus; Xl, X. laevis; Gg, G. gallus; Dr, D. rerio.*

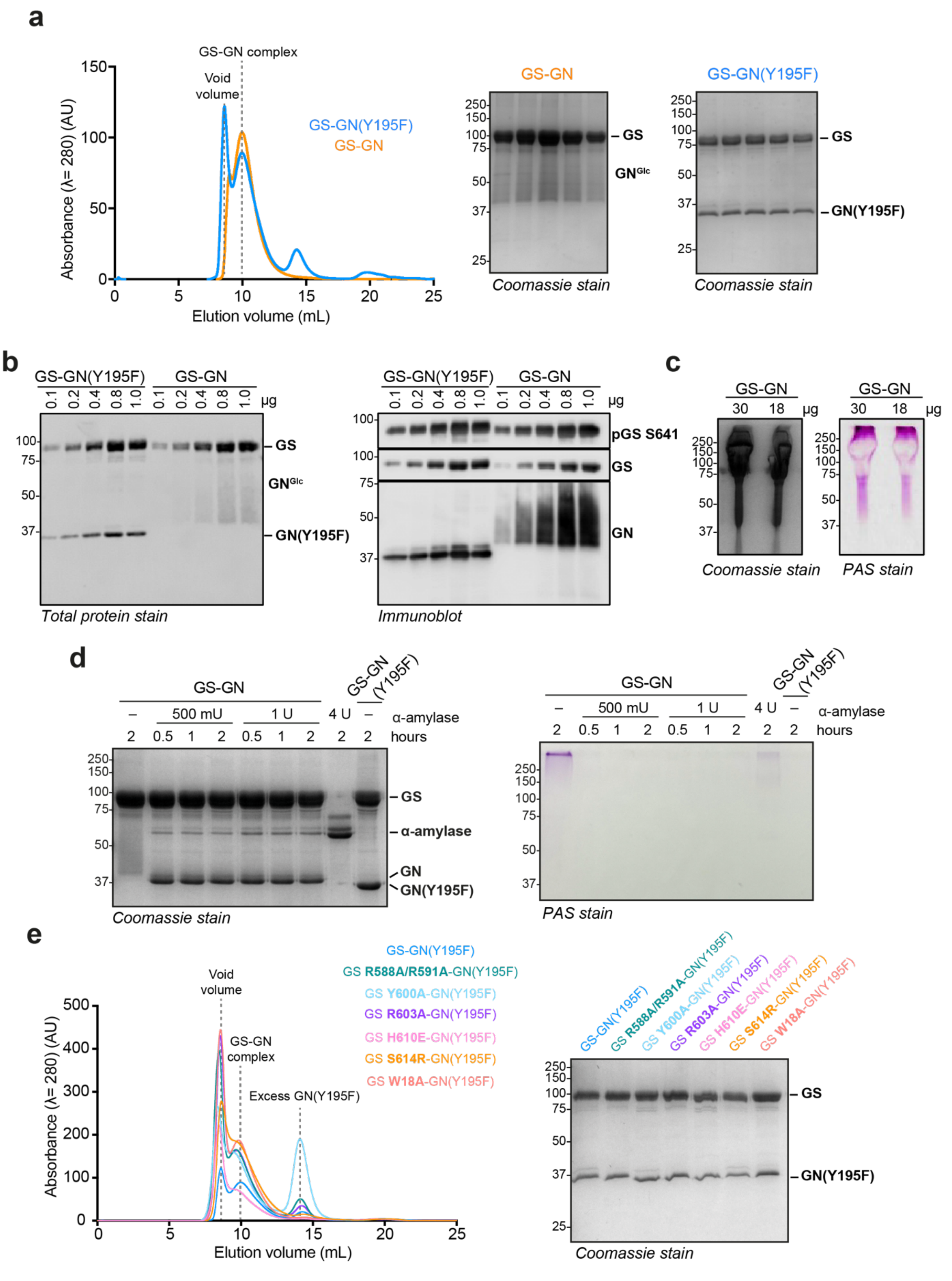

**Supplementary Fig. 3 Complex purifications and analysis**

**a** Size-exclusion chromatography profile of GS-GN Y195F (blue) and GS-GN WT (orange) (left), and SDS-PAGE analysis of the complex peak fractions (right). **b** Total protein stain and Immunoblot of GS-GN and GS-GN(Y195F) complexes. **c** Coomassie and PAS stain of GS-GN to show sugars associated with the WT complex in the smeared band. **d** Disappearance of the smeared band due to glucosylated GN, and appearance of single band for GN upon $\alpha$-amylase treatment. **e** Size-exclusion chromatography profile of GS WT and mutants in the GS-GN Y195F complex (left), and SDS-PAGE analysis of the corresponding mutants (right).

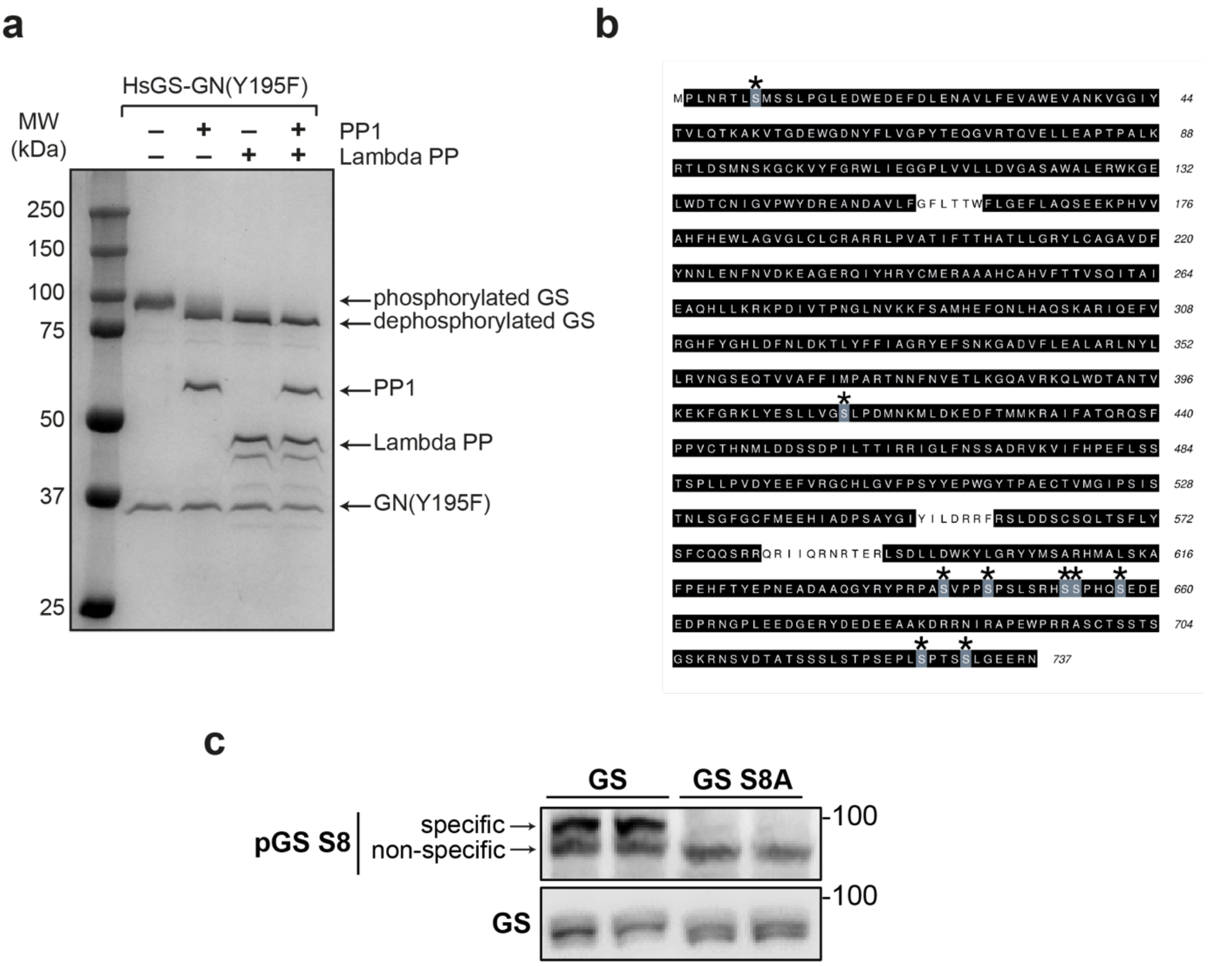

**Supplementary Fig. 4 Human GS produced in insect cells is phosphorylated**
**a** Shift in Hs GS band on SDS-PAGE gel to a lower molecular weight upon dephosphorylation by protein phosphatase 1 (PP1) and lambda protein phosphatase (lambda PP). **b** Sequence of human GS. Sequence coverage (96.74%) during phosphorylation site mapping is highlighted. Identified phosphorylated residues identified are highlighted in grey and annotated with an asterix. **c** Validation of the pGS S8 antibody.

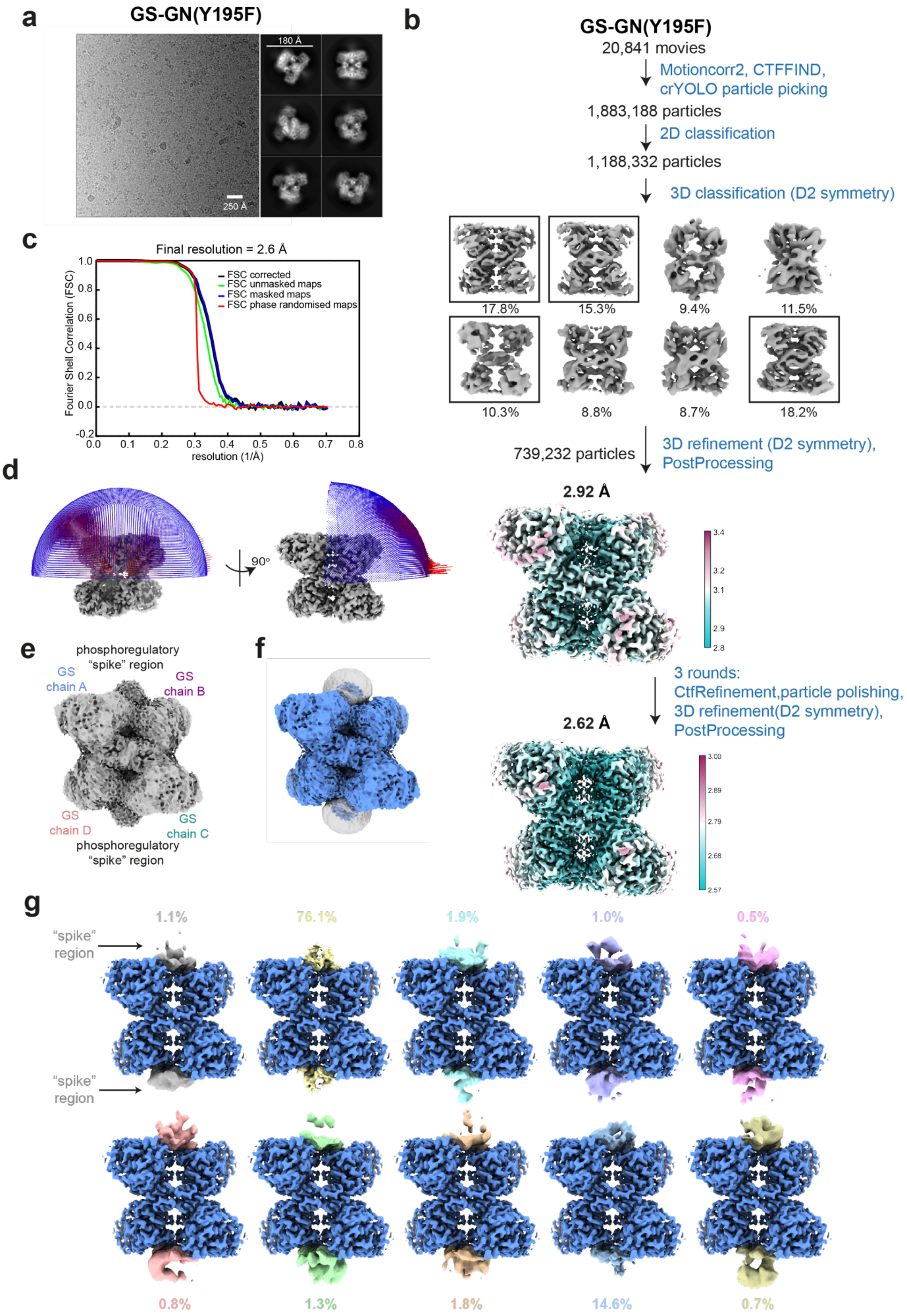
a
GS-GN(Y195F)
180 Å
250 Å
b
GS-GN(Y195F)
20,841 movies
Motioncorr2, CTFFIND, crYOLO particle picking
1,883,188 particles
2D classification
1,188,332 particles
3D classification (D2 symmetry)
17.8%
15.3%
9.4%
11.5%
10.3%
8.8%
8.7%
18.2%
739,232 particles
3D refinement (D2 symmetry), PostProcessing
2.92 Å
3.4
3.2
3.1
2.9
2.8
3 rounds: CtfRefinement,particle polishing, 3D refinement(D2 symmetry), PostProcessing
2.62 Å
3.00
2.89
2.79
2.68
2.57
c
Final resolution = 2.6 Å
FSC corrected
FSC unmasked maps
FSC masked maps
FSC phase randomised maps
Fourier Shell Correlation (FSC)
resolution (1/Å)
d
90°
e
phosphoregulatory "spike" region
GS chain A
GS chain B
GS chain D
GS chain C
phosphoregulatory "spike" region
f
g
"spike" region
"spike" region
1.1%
76.1%
1.9%
1.0%
0.5%
0.8%
1.3%
1.8%
14.6%
0.7%

**Supplementary Fig. 5 Cryo-electron microscopy analysis of human GS-GN Y195F complex**
**a** Representative micrograph, and a selection of 2D class averages. **b** Flow chart of data processing strategy. 3D classes boxed were selected for subsequent processing. Final map at a global resolution of 2.6 Å is coloured by local resolution. **c** FSC curve for the final map. The resolution was calculated using the gold-standard FSC cut-off at 0.143 frequency. **d** Euler angle distribution of particles used for the 3D reconstruction. Height of rods represents number of particles. **e** A 3D refinement map shown at a lower threshold to visualise the" spike" density. **f** Input map and mask used in the focussed 3D classification shown in g. **g** Focussed 3D classification of the phosphoregulatory "spike" region reveals flexibility of this inter subunit domain. 3D classes and percentage population are indicated with different colours and the 3D refined map of the core region is coloured blue.

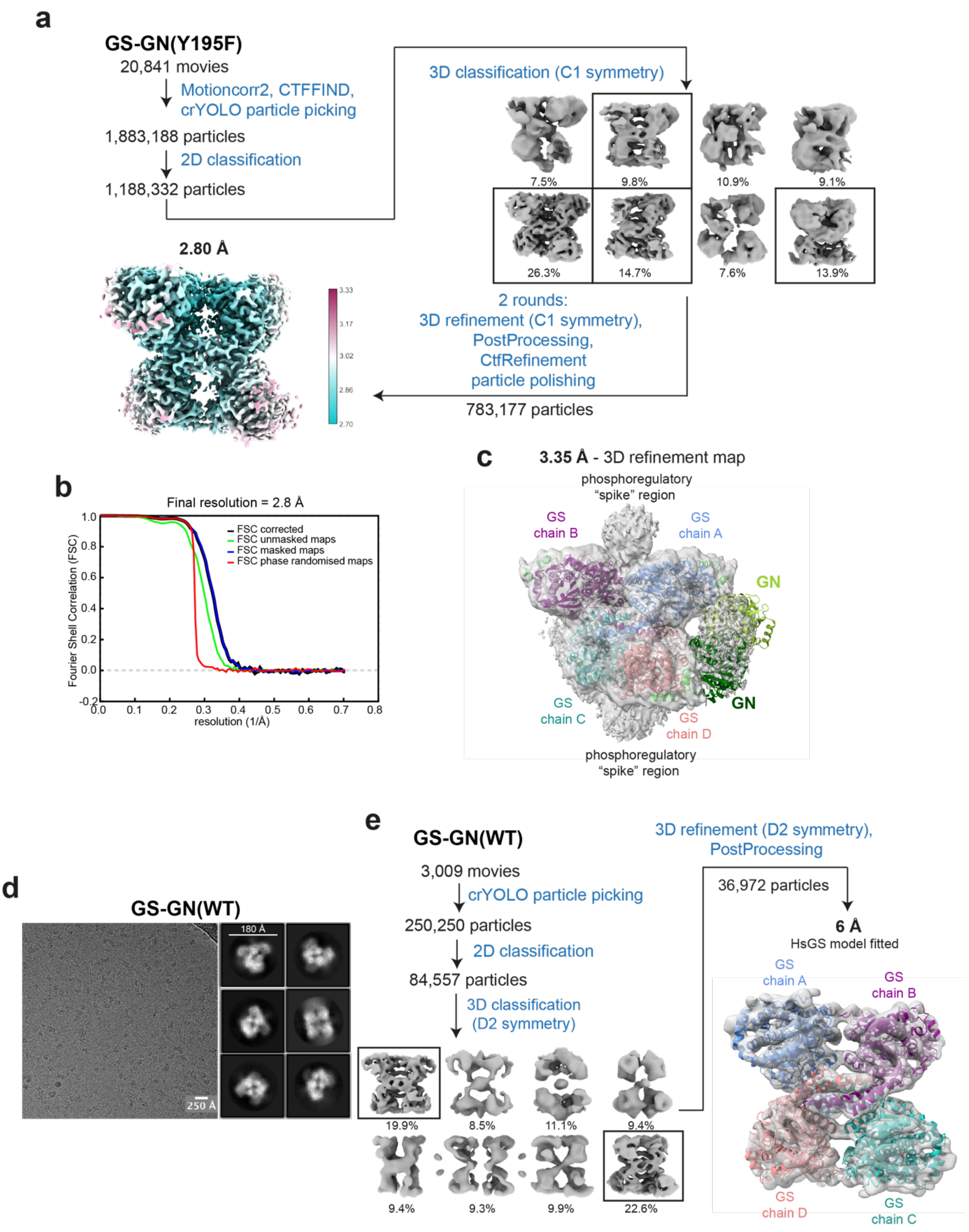

**Supplementary Fig. 6 Cryo-electron microscopy analysis of human GS-GN Y195F complex without the application of symmetry averaging and cryo-electron microscopy analysis of GS-GN**

**a** Flow chart of data processing strategy used. 3D classes boxed were selected for subsequent processing. Final map at a global resolution of 2.8 Å is shown. **b** FSC curve for the final map. The resolution was calculated using the gold-standard FSC cut-off at 0.143 frequency. **c** At a lower threshold, the 3.35 Å 3D refinement map has sufficient density for GN to allow the human GN crystal structure (PDB ID 3T7O) to be fitted. **d** Representative micrograph for the GS-GN cryo-EM dataset. **e** GS-GN WT 6 Å map fits HsGS structure, showing little structural difference between the GS-GN(WT) and GS-GN(Y195F) complexes, at this resolution.

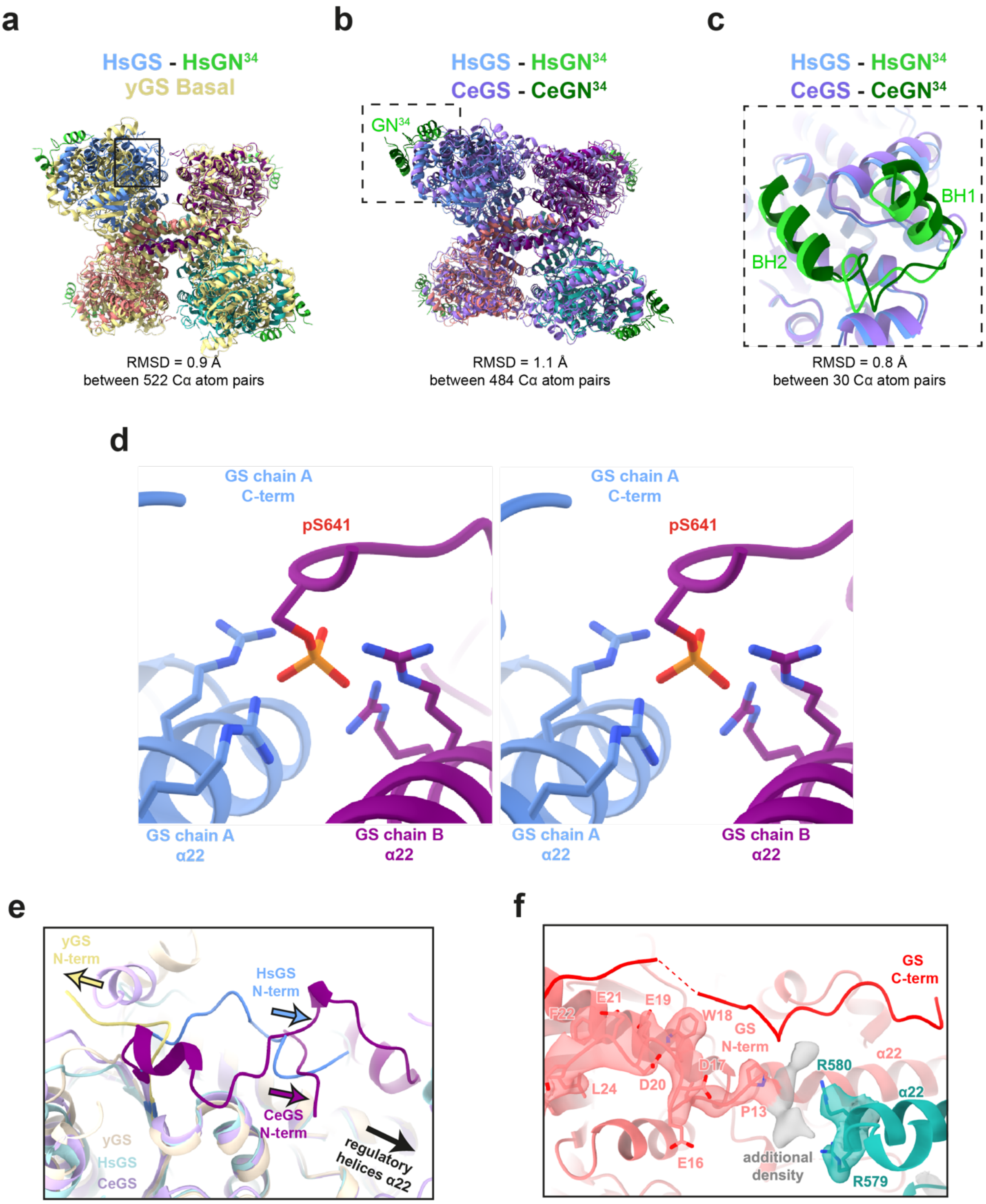

**Supplementary Fig. 7 Comparison of human tetrameric GS with yeast GS and *C. elegans* GS-GN$^{34}$ crystal structures**

**a** Human GS has a tetrameric arrangement, agreeing with crystal structures of yeast (yellow, PDB ID 3NAZ) **b** and *C. elegans* (PDB ID 4QLB). The area boxed is shown in **c** HsGN$^{34}$ is consistent with the CeGS-GN$^{34}$ structure (PDB ID 4QLB). **d** Stereoview of pS641 interacting with the arginine cradle on the regulatory helices α22. **e** Human and *C. elegans* N-terminal tails are situated towards the regulatory helices. However, yeast N-terminus is pointing away from the regulatory helices. **f** Additional density for the human N-terminal tail can be seen near the regulatory helices, potentially interacting with R579 and R580 on the adjacent protomer. The LAFTER denoised map using C1 symmetry is shown.

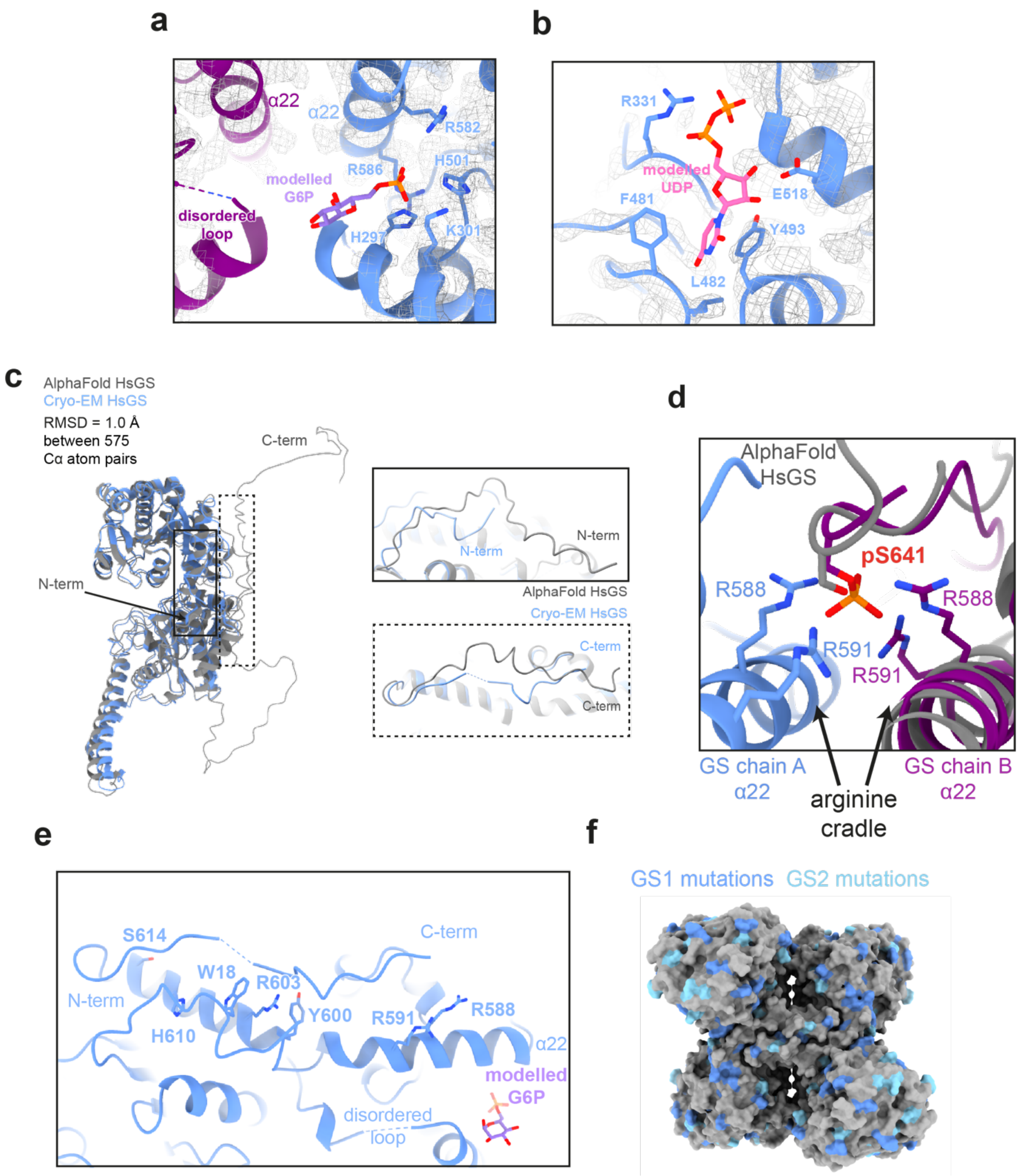

**Supplementary Fig. 8 Structural analysis of inhibited human GS**
**a** Close up view of the allosteric binding site, G6P is modelled using the yeast GS + G6P structure (PDB ID 5SUK). The residues involved in the G6P interaction are shown as sticks. **b** Close up view of the active site. UDP is modelled in the active site and the residues involved in the interaction are shown and labelled. **c** Comparison of human GS monomer model with the AlphaFold model (accessed on 1 October 2021) and position of the N- and C- terminal tails **d** Comparison of human GS pS641 model with the AlphaFold model. **e** The N- and C-terminus of one GS protomer. The residues shown are mutated in this study. G6P is modelled and the disordered loop is labelled. **f** Human GS1 and GS2 mutants mapped onto a surface representation of HsGS (reported here).

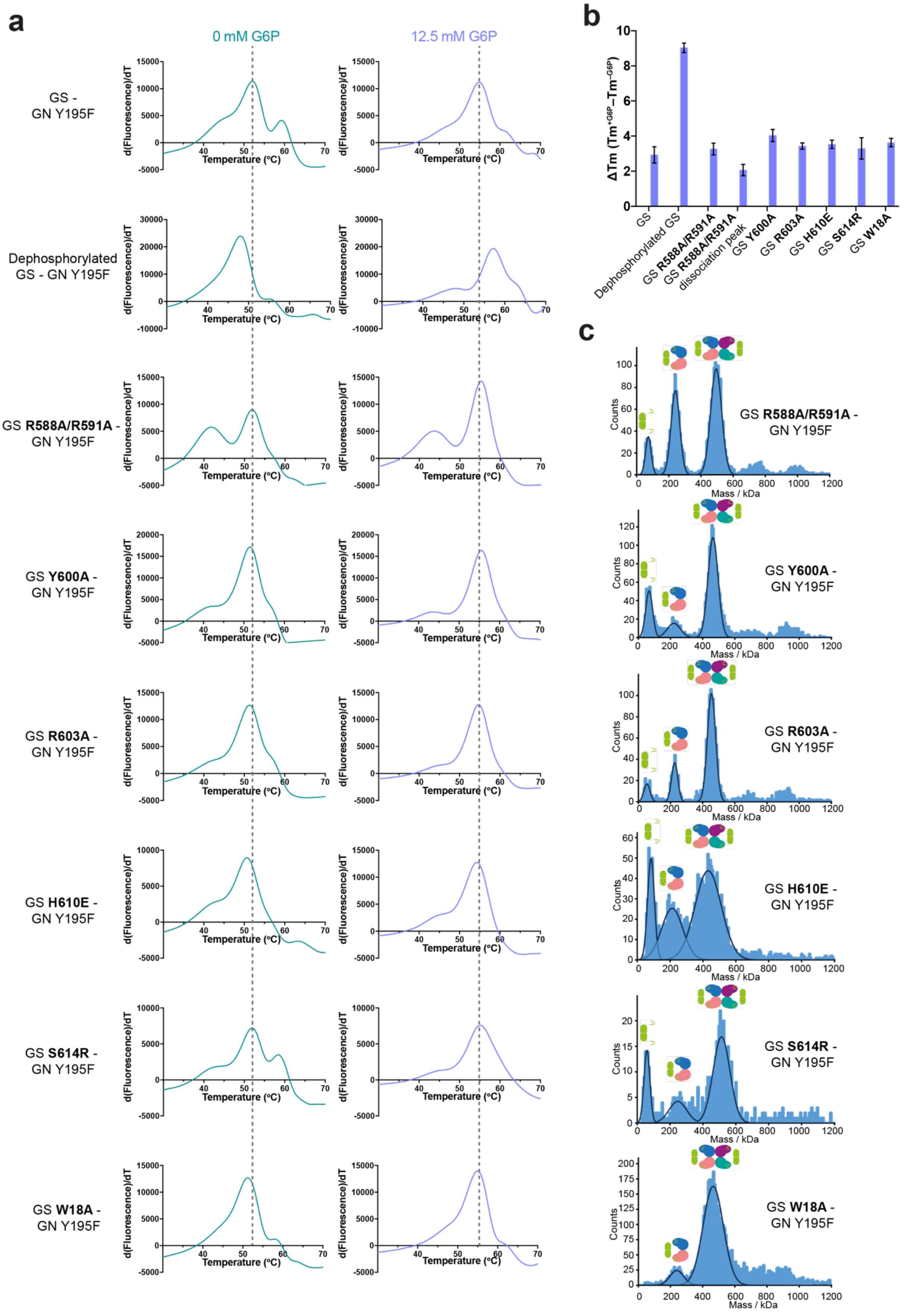
a
0 mM G6P
12.5 mM G6P
GS - GN Y195F
Dephosphorylated GS - GN Y195F
GS R588A/R591A - GN Y195F
GS Y600A - GN Y195F
GS R603A - GN Y195F
GS H610E - GN Y195F
GS S614R - GN Y195F
GS W18A - GN Y195F
d(Fluorescence)/dT
Temperature (ºC)
b
ΔTm (Tm+G6P–Tm−G6P)
GS
Dephosphorylated GS
GS R588A/R591A
GS R588A/R591A dissociation peak
GS Y600A
GS R603A
GS H610E
GS S614R
GS W18A
c
GS R588A/R591A - GN Y195F
GS Y600A - GN Y195F
GS R603A - GN Y195F
GS H610E - GN Y195F
GS S614R - GN Y195F
GS W18A - GN Y195F
Counts
Mass / kDa

**Supplementary Fig. 9 Stability of GS WT and mutants in the GS-GN(Y195F) complex**
**a** Differential scanning fluorimetry (DSF) analysis of GS WT, various mutants and dephosphorylated GS in the GS-GN(Y195F) complex. Addition of G6P leads to stabilisation of the complex, seen by an increase in melting temperature **b** Changes in melting temperature (ΔTm) was calculated by subtracting the Tm of 0 mM G6P of the corresponding protein. Data in **a** and **b** are mean from n = 3 experiments carried out in technical duplicates (dephosphorylated GS) and triplicates (WT and mutant GS). **c** Mass photometry analysis of GS mutants in the GS-GN(Y195F).

**Supplementary Table 1:** Cryo-EM data collection, refinement and validation statistics

| | **GS-GN(Y195F) PDB 7ZBN EMDB-14587** | **GS-GN** |
|---|---|---|
| **Data collection and processing** | | |
| Microscope | FEI Titan KRIOS | FEI Titan KRIOS |
| Detector | Falcon 4 | Falcon 3 |
| Energy Filter (eV) | 10 | / |
| Magnification (x) | 165,000 | 75,000 |
| Voltage (kV) | 300 | 300 |
| Electron exposure ($e^-/Å^2$) | 34.8 | 85 |
| Defocus range (μm) | -1 to -2.2 | -1.7 to -3.1 |
| Pixel size (Å) | 0.71 | 1.065 |
| Initial particle images (no.) | 1,883,188 | 250,250 |
| Final particle images, D2 symmetry (no.) | 739,232 | 36,972 |
| Final particle images, C1 symmetry (no.) | 783,177 | / |
| Map resolution, D2 symmetry (Å) | 2.62 | 6 |
| Map resolution, C1 symmetry (Å) | 2.8 | / |
| FSC threshold | 0.143 | 0.143 |
| **Refinement** | | |
| Model resolution (Å) | 2.7 | |
| FSC threshold | 0.5 | |
| *Model composition* | | |
| Non-hydrogen atoms | 41588 | |
| Protein residues | 2608 | |
| Ligands | 0 | |
| *B factors ($Å^2$)* | | |
| Protein (min/max/mean) | 5.81/46.06/19.42 | |
| *Map-model CC* | | |
| CC (mask) | 0.81 | |
| CC (volume) | 0.76 | |
| CC (peaks) | 0.69 | |
| CC (box) | 0.71 | |
| *R.m.s deviations* | | |
| Bond lengths (Å) | 0.006 | |
| Bond angle (°) | 0.551 | |
| **Validation** | | |
| Molprobity score | 1.71 | |
| Clashscore | 10.45 | |
| Rotamers outliers (%) | 0 | |
| *Ramachandran plot* | | |
| Favored (%) | 97.01 | |
| Allowed (%) | 2.99 | |
| Outliers (%) | 0 | |

**Supplementary Table 2:** Mutations of GS1 and GS2 genes. Source: clinvar, last accessed 4 October 2021

| GS1 mutations<br>CDS mutation | AA mutation | Type | Accession code (clinvar) | Condition(s) | Residue interaction |
|---|---|---|---|---|---|
| c.1A>C | M1L | missense | VCV000430320 | Glycogen storage disease 0, muscle | |
| c.3G>T | M1I | missense | VCV001039847 | Glycogen storage disease 0, muscle | |
| c.20T>C | L7S | missense | VCV000999389 | Glycogen storage disease 0, muscle | |
| c.40G>C | G14R | missense | VCV000964057 | Glycogen storage disease 0, muscle | |
| c.56A>T | E19V | missense | VCV001063670 | not provided | |
| c.80C>T | A27V | missense | VCV001064089 | Glycogen storage disease 0, muscle | |
| c.101G>T | W34L | missense | VCV000429776 | Glycogen storage disease 0, muscle | |
| c.114C>G | N38K | missense | VCV000946451 | Glycogen storage disease 0, muscle | |
| c.134C>T | T45M | missense | VCV000945011 | Glycogen storage disease 0, muscle | |
| c.178G>A | D60N | missense | VCV001002555 | Glycogen storage disease 0, muscle | |
| c.213G>C | Q71H | missense | VCV000642436 | not provided | |
| c.214G>T | G72C | missense | VCV001014411 | Glycogen storage disease 0, muscle | |
| c.217G>A | V73M | missense | VCV000970915 | Glycogen storage disease 0, muscle | |
| c.221G>C | R74T | missense | VCV000645191 | Glycogen storage disease 0, muscle | |
| c.254C>T | P85L | missense | VCV000960357 | Glycogen storage disease 0, muscle | |
| c.278C>T | S93F | missense | VCV000854343 | Glycogen storage disease 0, muscle | |
| c.310G>A | G104R | missense | VCV000450024 | not provided | |
| c.314G>A | R105H | missense | VCV000654802 | Glycogen storage disease 0, muscle | |
| c.324C>G | I108M | missense | VCV000998960 | Glycogen storage disease 0, muscle | |
| c.362C>T | A121V | missense | VCV001040534 | Glycogen storage disease 0, muscle | |
| c.395A>G | E132G | missense | VCV000567434 | Glycogen storage disease 0, muscle | |
| c.418G>A | G140R | missense | VCV001004862 | Glycogen storage disease 0, muscle | GN[34] interaction |
| c.425C>T | P142L | missense | VCV000941325 | Glycogen storage disease 0, muscle | GN[34] interaction |
| c.448G>A | D150N | missense | VCV000942370 | Glycogen storage disease 0, muscle | |
| c.500C>A | A167E | missense | VCV000652496 | Glycogen storage disease 0, muscle | |
| c.505A>G | S169G | missense | VCV000894589 | Glycogen storage disease 0, muscle | |
| c.524T>C | V175A | missense | VCV000946427 | Glycogen storage disease 0, muscle | |
| c.553G>A | G185S | missense | VCV000862647 | Glycogen storage disease 0, muscle | |
| c.556G>A | V186I | missense | VCV000329823 | Glycogen storage disease 0, muscle | |
| c.578C>T | A193V | missense | VCV000847954 | Glycogen storage disease 0, muscle | GN[34] interaction |
| c.584G>A | R195Q | missense | VCV000967367 | Glycogen storage disease 0, muscle | GN[34] interaction |
| c.631C>T | R211C | missense | VCV000970295 | Glycogen storage disease 0, muscle | |
| c.646G>A | G216S | missense | VCV000939691 | Glycogen storage disease 0, muscle | |
| c.646G>T | G216C | missense | VCV000966878 | Glycogen storage disease 0, muscle | |
| c.650C>T | A217V | missense | VCV000949974 | Glycogen storage disease 0, muscle | |
| c.652G>A | V218M | missense | VCV000835130 | Glycogen storage disease 0, muscle | |

| | | | | | |
|---|---|---|---|---|---|
| c.652G>C | V218L | missense | VCV001054282 | Glycogen storage disease 0, muscle | |
| c.666C>G | N222K | missense | VCV000836150 | Glycogen storage disease 0, muscle | sugar binding |
| c.666C>A | N222K | missense | VCV001037659 | Glycogen storage disease 0, muscle | sugar binding |
| c.683A>G | N228S | missense | VCV001007054 | Glycogen storage disease 0, muscle | GN[34] interaction |
| c.704A>T | E235V | missense | VCV000856687 | Glycogen storage disease 0, muscle | |
| c.722G>A | R241Q | missense | VCV001018886 | Glycogen storage disease 0, muscle | |
| c.754G>A | A252T | missense | VCV000391854 | Glycogen storage disease 0, muscle | |
| c.760G>A | V254I | missense | VCV000940671 | Glycogen storage disease 0, muscle | |
| c.793G>A | E265K | missense | VCV000653921 | Glycogen storage disease 0, muscle | |
| c.803A>T | H268L | missense | VCV000894180 | Glycogen storage disease 0, muscle | |
| c.872A>G | H291R | missense | VCV001023691 | Glycogen storage disease 0, muscle | G6P binding |
| c.944A>T | H315L | missense | VCV001015336 | Glycogen storage disease 0, muscle | |
| c.1012G>C | G338R | missense | VCV000945364 | Glycogen storage disease 0, muscle | |
| c.1015G>T | A339S | missense | VCV001013781 | Glycogen storage disease 0, muscle | |
| c.1020C>A | D340E | missense | VCV000955642 | Glycogen storage disease 0, muscle | |
| c.1021G>A | V341I | missense | VCV000329819 | Glycogen storage disease 0, muscle | |
| c.1049A>G | N350S | missense | VCV001024950 | Glycogen storage disease 0, muscle | |
| c.1052A>G | Y351C | missense | VCV000659273 | Glycogen storage disease 0, muscle | sugar binding |
| c.1075G>A | E359K | missense | VCV001056519 | Glycogen storage disease 0, muscle | |
| c.1077_1078delinsAA | Q360K | missense | VCV000858591 | Glycogen storage disease 0, muscle | |
| c.1078C>A | Q360K | missense | VCV000894178 | Glycogen storage disease 0, muscle | |
| c.1112G>A | R371Q | missense | VCV000844943 | Glycogen storage disease 0, muscle | |
| c.1121A>G | N374S | missense | VCV000585159 | Glycogen storage disease 0, muscle | |
| c.1127A>G | N376S | missense | VCV001007393 | Glycogen storage disease 0, muscle | sugar binding |
| c.1144G>A | G382S | missense | VCV000944904 | Glycogen storage disease 0, muscle | |
| c.1145G>A | G382D | missense | VCV000329816 | Glycogen storage disease 0, muscle | tetramerization domain |
| c.1156C>T | R386C | missense | VCV000893350 | Glycogen storage disease 0, muscle | tetramerization domain |
| c.1184C>T | T395M | missense | VCV000893348 | Glycogen storage disease 0, muscle | tetramerization domain |
| c.1190A>G | K397R | missense | VCV000641050 | Glycogen storage disease 0, muscle | tetramerization domain |
| c.1236C>G | S412R | missense | VCV000662709 | Glycogen storage disease 0, muscle | phosphorylation site (identified by phosphorylation mapping) |
| c.1243G>A | D415N | missense | VCV000808611 | Glycogen storage disease 0, muscle | |
| c.1246A>G | M416V | missense | VCV000329814 | Glycogen storage disease 0, muscle | tetramerization domain |
| c.1279A>G | M427V | missense | VCV001017549 | Glycogen storage disease 0, muscle | |
| c.1303A>G | T435A | missense | VCV000893132 | Glycogen storage disease 0, muscle | |

| | | | | | |
|---|---|---|---|---|---|
| c.1315_1316delinsAA | S439N | missense - | VCV000649510 | Glycogen storage disease 0, muscle | |
| c.1318T>C | F440L | missense | VCV000498163 | not provided | sugar binding |
| c.1324C>G | P442A | missense | VCV000498167 | Glycogen storage disease 0, muscle | sugar binding |
| c.1324C>A | P442T | missense | VCV000936512 | Glycogen storage disease 0, muscle | sugar binding |
| c.1363C>T | P455S | missense | VCV001004987 | Glycogen storage disease 0, muscle | |
| c.1407T>A | S469R | missense | VCV000893129 | Glycogen storage disease 0, muscle | |
| c.1412A>G | D471G | missense | VCV000947297 | Glycogen storage disease 0, muscle | |
| c.1432C>T | H478Y | missense | VCV000942790 | Glycogen storage disease 0, muscle | |
| c.1436C>T | P479L | missense | VCV000643030 | Glycogen storage disease 0, muscle | |
| c.1475A>T | D492V | missense | VCV000938769 | Glycogen storage disease 0, muscle | |
| c.1492C>T | R498C | missense | VCV001014236 | Glycogen storage disease 0, muscle | |
| c.1520C>T | S507F | missense | VCV000546850 | not provided | |
| c.1559C>T | T520M | missense | VCV001026554 | Glycogen storage disease 0, muscle | |
| c.1615G>A | E539K | missense | VCV000329810 | Neuroferritinopathy, Hereditary hyperferritinemia with congenital cataracts, Glycogen storage disease 0, muscle | sugar binding |
| c.1754A>G | Q585R | missense | VCV000859569 | Glycogen storage disease 0, muscle | regulatory helix, α23 |
| c.1822G>A | A608T | missense | VCV001016164 | Glycogen storage disease 0, muscle | |
| c.1823C>G | A608G | missense | VCV001052744 | Glycogen storage disease 0, muscle | |
| c.1834G>C | A612P | missense | VCV001051757 | Glycogen storage disease 0, muscle | |
| c.1835C>T | A612V | missense | VCV001061438 | Glycogen storage disease 0, muscle | |
| c.1859A>G | H620R | missense | VCV001043805 | Glycogen storage disease 0, muscle | |
| c.1879G>A | E627K | missense | VCV000576955 | Glycogen storage disease 0, muscle | |
| c.1898G>T | G633V | missense | VCV001021949 | Glycogen storage disease 0, muscle | |
| c.1909C>G | P637A | missense | VCV000651479 | Glycogen storage disease 0, muscle | |
| c.1922C>T | S641L | missense | VCV000916245 | Glycogen storage disease 0, muscle | phosphorylation site |
| c.1961C>T | P654L | missense | VCV000329807 | Glycogen storage disease 0, muscle | |
| c.1985A>G | D662G | missense | VCV000934446 | Glycogen storage disease 0, muscle | |
| c.1990C>T | R664W | missense | VCV000329806 | Glycogen storage disease 0, muscle | |
| c.2008G>A | E670K | missense | VCV001009397 | Glycogen storage disease 0, muscle | |
| c.2009A>G | E670G | missense | VCV000999215 | Glycogen storage disease 0, muscle | |
| c.2014G>A | G672S | missense | VCV000945494 | Glycogen storage disease 0, muscle | |
| c.2017G>A | E673K | missense | VCV000957665 | Glycogen storage disease 0, muscle | |
| c.2021G>A | R674H | missense | VCV001021262 | Glycogen storage disease 0, muscle | |
| c.2032G>A | D678N | missense | VCV001024883 | Glycogen storage disease 0, muscle | |
| c.2035G>A | E679K | missense | VCV000894150 | Glycogen storage disease 0, muscle | |
| c.2042C>T | A681V | missense | VCV000985784 | Inborn genetic diseases | |

| | | | | | |
|---|---|---|---|---|---|
| c.2044G>A | A682T | missense | VCV001018906 | Glycogen storage disease 0, muscle | |
| c.2065C>T | R689C | missense | VCV001006088 | Glycogen storage disease 0, muscle | |
| c.2096G>T | C699F | missense | VCV001010299 | Glycogen storage disease 0, muscle | |
| c.2105C>T | S702F | missense | VCV000836504 | Glycogen storage disease 0, muscle | |
| c.2120A>C | K707T | missense | VCV000894149 | Glycogen storage disease 0, muscle | |
| c.2122C>G | R708G | missense | VCV000844469 | Glycogen storage disease 0, muscle | |
| c.2206C>T | R736C | missense | VCV000837003 | Glycogen storage disease 0, muscle | |
| c.2207G>A | R736H | missense | VCV000329804 | Glycogen storage disease 0, muscle, Neuroferritinopathy, Hereditary hyperferritinemia with congenital cataracts | |
| c.907C>T | R303* | nonsense | VCV000965769 | Glycogen storage disease 0, muscle | |
| c.913C>T | Q305* | nonsense | VCV001074189 | Glycogen storage disease 0, muscle | |
| c.1384C>T | R462* | nonsense | VCV000016057 | Glycogen storage disease 0, muscle | sugar binding |
| c.1615G>T | E539* | nonsense | VCV000570876 | Glycogen storage disease 0, muscle | sugar binding |
| c.160dup | T54fs | frameshift - duplication | VCV000422359 | not provided | |
| c.162_163del | D56fs | frameshift - deletion | VCV000128236 | Glycogen storage disease 0, muscle | |
| c.699_700del | R236fs | frameshift - deletion | VCV001069149 | Glycogen storage disease 0, muscle | |
| c.1204del | R402fs | frameshift - deletion | VCV000567037 | Glycogen storage disease 0, muscle | |
| c.2207del | R736fs | frameshift - deletion | VCV000861694 | Glycogen storage disease 0, muscle | |

**GS2 mutations**

| CDS mutation | AA mutation | Type | Accession code (clinvar) | Condition(s) | Residue interaction |
|---|---|---|---|---|---|
| c.50A>G | Q17R | missense | VCV000665936 | Glycogen storage disease due to hepatic glycogen synthase deficiency | |
| c.116A>G | N39S | missense | VCV000016054 | Glycogen storage disease due to hepatic glycogen synthase deficiency | |
| c.154G>A | A52T | missense | VCV000308011 | Glycogen storage disease due to hepatic glycogen synthase deficiency | |
| c.215A>G | H72R | missense | VCV000884167 | Glycogen storage disease due to hepatic glycogen synthase deficiency | |
| c.279C>G | D93E | missense | VCV000214522 | not specified | |
| c.280G>A | A94T | missense | VCV000137522 | Glycogen storage disease due to hepatic glycogen synthase deficiency | |
| c.289A>G | K97E | missense | VCV000308009 | Glycogen storage disease due to hepatic glycogen synthase deficiency | sugar binding |
| c.299G>A | C100Y | missense | VCV000214527 | not provided | |
| c.395G>A | G132D | missense | VCV000214523 | not specified | |
| c.421G>A | G141S | missense | VCV000137523 | Glycogen storage disease due to hepatic glycogen synthase deficiency | |
| c.427C>T | P143S | missense | VCV000861394 | Glycogen storage disease due to hepatic glycogen synthase deficiency | GN[34] interaction |

| | | | | | |
|---|---|---|---|---|---|
| c.470C>T | S157F | missense | VCV000999043 | Glycogen storage disease due to hepatic glycogen synthase deficiency | |
| c.520T>C | Y174H | missense | VCV000380333 | not specified | |
| c.526G>A | V176I | missense | VCV000736468 | Glycogen storage disease due to hepatic glycogen synthase deficiency | |
| c.556A>T | I186F | missense | VCV000214524 | not specified | |
| c.577G>A | A193T | missense | VCV000261471 | Glycogen storage disease due to hepatic glycogen synthase deficiency | GN[34] interaction |
| c.653T>C | I218T | missense | VCV000964160 | Glycogen storage disease due to hepatic glycogen synthase deficiency | |
| c.753C>G | C251W | missense | VCV001036478 | Glycogen storage disease due to hepatic glycogen synthase deficiency | GN[34] interaction |
| c.755C>G | A252G | missense | VCV000308003 | Glycogen storage disease due to hepatic glycogen synthase deficiency | |
| c.799G>A | E267K | missense | VCV000882213 | Glycogen storage disease due to hepatic glycogen synthase deficiency | |
| c.956A>T | D319V | missense | VCV000376836 | not provided | |
| c.1015G>C | A339P | missense | VCV000016052 | Glycogen storage disease due to hepatic glycogen synthase deficiency | |
| c.1087A>G | M363V | missense | VCV000261462 | Glycogen storage disease due to hepatic glycogen synthase deficiency | |
| c.1129G>A | V377M | missense | VCV000880824 | Glycogen storage disease due to hepatic glycogen synthase deficiency | |
| c.1171G>T | D391Y | missense | VCV000884120 | Glycogen storage disease due to hepatic glycogen synthase deficiency | tetramerization domain |
| c.1245C>G | D415E | missense | VCV000137524 | Glycogen storage disease due to hepatic glycogen synthase deficiency | tetramerization domain |
| c.1277C>T | T426I | missense | VCV000214531 | not provided | tetramerization domain |
| c.1279A>C | I427L | missense | VCV001058720 | Glycogen storage disease due to hepatic glycogen synthase deficiency | |
| c.1334C>T | T445M | missense | VCV000952317 | Glycogen storage disease due to hepatic glycogen synthase deficiency | |
| c.1336C>G | H446D | missense | VCV000016056 | Glycogen storage disease due to hepatic glycogen synthase deficiency | |
| c.1418T>G | V473G | missense | VCV000307993 | Glycogen storage disease due to hepatic glycogen synthase deficiency | |
| c.1427T>A | I476N | missense | VCV000214532 | not provided | |
| c.1436C>A | P479Q | missense | VCV000016051 | Glycogen storage disease due to hepatic glycogen synthase deficiency | |
| c.1447T>C | S483P | missense | VCV000016055 | Glycogen storage disease due to hepatic glycogen synthase deficiency | |
| c.1472T>G | M491R | missense | VCV000016053 | Glycogen storage disease due to hepatic glycogen synthase deficiency | |
| c.1475A>T | D492V | missense | VCV000418239 | not provided | |
| c.1477T>A | Y493N | missense | VCV000521873 | Inborn genetic diseases | active site |
| c.1522T>G | Y508D | missense | VCV000418240 | not provided | sugar binding |
| c.1549G>C | A517P | missense | VCV000307992 | Glycogen storage disease due to hepatic glycogen synthase deficiency | |
| c.1553A>C | E518A | missense | VCV000214533 | Glycogen storage disease due to hepatic glycogen synthase deficiency | active site |

| | | | | | |
|---|---|---|---|---|---|
| c.1636A>G | T546A | missense | VCV000137526 | Glycogen storage disease due to hepatic glycogen synthase deficiency | |
| c.1657G>A | V553I | missense | VCV000214525 | not specified | |
| c.1672C>T | R558C | missense | VCV000648650 | Glycogen storage disease due to hepatic glycogen synthase deficiency | sugar binding |
| c.1745G>A | R582K | missense | VCV000534624 | Glycogen storage disease due to hepatic glycogen synthase deficiency | G6P binding |
| c.1774C>G | L592V | missense | VCV000653240 | Glycogen storage disease due to hepatic glycogen synthase deficiency | regulatory helix, α23 |
| c.1789G>A | D597N | missense | VCV000214535 | not provided | |
| c.1790A>G | D597G | missense | VCV000307990 | Glycogen storage disease due to hepatic glycogen synthase deficiency | |
| c.1820A>G | H607R | missense | VCV000307989 | Glycogen storage disease due to hepatic glycogen synthase deficiency | |
| c.1829A>G | H610R | missense | VCV000936980 | Glycogen storage disease due to hepatic glycogen synthase deficiency | |
| c.1880C>T | S627T | missense | VCV000766741 | Glycogen storage disease due to hepatic glycogen synthase deficiency | |
| c.1889C>T | T630M | missense | VCV000719047 | Glycogen storage disease due to hepatic glycogen synthase deficiency | |
| c.1906T>C | Y636H | missense | VCV000658693 | Glycogen storage disease due to hepatic glycogen synthase deficiency | |
| c.1965G>C | Q655H | missense | VCV000137528 | Glycogen storage disease due to hepatic glycogen synthase deficiency | |
| c.2005G>A | D669N | missense | VCV000214526 | Glycogen storage disease due to hepatic glycogen synthase deficiency | |
| c.2054T>C | F685S | missense | VCV000137529 | Glycogen storage disease due to hepatic glycogen synthase deficiency, not provided | |
| c.2068G>T | V690F | missense | VCV000720229 | not provided | |
| c.2072C>T | P691L | missense | VCV000798247 | Glycogen storage disease due to hepatic glycogen synthase deficiency | |
| c.547C>T | Q183* | nonsense | VCV000214529 | Glycogen storage disease due to hepatic glycogen synthase deficiency | |
| c.574C>T | R192* | nonsense | VCV000214530 | Glycogen storage disease due to hepatic glycogen synthase deficiency | GN[34] interaction |
| c.736C>T | R246* | nonsense | VCV000016049 | Glycogen storage disease due to hepatic glycogen synthase deficiency | GN[34] interaction |
| c.925C>T | R309* | nonsense | VCV000078954 | Glycogen storage disease due to hepatic glycogen synthase deficiency | |
| c.1156C>T | R386* | nonsense | VCV000569452 | Glycogen storage disease due to hepatic glycogen synthase deficiency | |
| c.465del | F155fs | frameshift - deletion | VCV000937499 | Glycogen storage disease due to hepatic glycogen synthase deficiency | |
| c.1081del | T361fs | frameshift - deletion | VCV000667422 | Glycogen storage disease | |
| c.1974dup | V659fs | frameshift - duplication | VCV000631679 | Glycogen storage disease due to hepatic glycogen synthase deficiency | |
